# The transcriptome of the avian malaria parasite *Plasmodium ashfordi* displays host-specific gene expression

**DOI:** 10.1101/072454

**Authors:** Elin Videvall, Charlie K. Cornwallis, Dag Ahrén, Vaidas Palinauskas, Gediminas Valkiūnas, Olof Hellgren

## Abstract

Malaria parasites (*Plasmodium* spp.) include some of the world’s most widespread and virulent pathogens. Our knowledge of the molecular mechanisms these parasites use to invade and exploit hosts other than mice and primates is, however, extremely limited. It is therefore imperative to characterize transcriptome-wide gene expression from non-model malaria parasites and how this varies across host individuals. Here, we used high-throughput Illumina RNA-sequencing on blood from wild-caught Eurasian siskins experimentally infected with a clonal strain of the avian malaria parasite *Plasmodium ashfordi* (lineage GRW2). By using a multi-step approach to filter out host transcripts, we successfully assembled the blood-stage transcriptome of *P. ashfordi.* A total of 11 954 expressed transcripts were identified, and 7 860 were annotated with protein information. We quantified gene expression levels of all parasite transcripts across three hosts during two infection stages – peak and decreasing parasitemia. Interestingly, parasites from the same host displayed remarkably similar expression profiles during different infection stages, but showed large differences across hosts, indicating that *P. ashfordi* may adjust its gene expression to specific host individuals. We further show that the majority of transcripts are most similar to the human parasite *Plasmodium falciparum,* and a large number of red blood cell invasion genes were discovered, suggesting evolutionary conserved invasion strategies between mammalian and avian *Plasmodium.* The transcriptome of *P. ashfordi* and its host-specific gene expression advances our understanding of *Plasmodium* plasticity and is a valuable resource as it allows for further studies analysing gene evolution and comparisons of parasite gene expression.

## Introduction

The apicomplexan parasites of the genus *Plasmodium* (malaria parasites) encompass a worldwide distribution and infect a multitude of vertebrate hosts, including reptiles, birds and mammals (Garnham 1966). Their virulence can be highly variable between different strains and species. Some induce mild pathogenic effects on hosts and some cause severe disease, leading to high mortality rates (Palinauskas *et al.* 2008). Host individuals and host species also differ in their resistance and tolerance to malaria, and this interaction between host and parasite ultimately determines disease severity. Furthermore, the molecular response of hosts changes during the course of infection and creates a dynamic environment in which the parasites need to accommodate. Nevertheless, our understanding of how malaria parasites respond molecularly to different host individuals and to changes in the host immune defence over time is very limited.

Parasites of two clades in *Plasmodium* have been extensively studied from a molecular perspective, murine and primate parasites. We have learned a great deal about how malaria parasites of humans evolved and function by studying transcriptomes of their rodent-infecting relatives (Hall *et al.* 2005; Spence *et al.* 2013; Otto *et al.* 2014a). The majority of studies investigating gene expression in human malaria parasites have been conducted using cell lines (*in vitro*) or tissue cultures (*ex vivo*), which has provided tremendous insight into the biology of *Plasmodium* life stages (see e.g. Bozdech *et al.* 2003; Otto *et al.* 2010; Siegel *et al.* 2014). However, several discrepancies in parasite expression between cultures and live animals (*in vivo*) have been documented (Lapp *et al.* 2015) and a wide range of host environmental factors are absent in the *in vitro* systems. For example, temperature fluctuations, inflammatory and immune effector molecules, hormones, metabolites, microenvironments, and varying levels of oxygen, pH, and glucose are difficult to simulate in *in vitro* settings (LeRoux *et al.* 2009). Parasites cultured outside hosts reflect this with different expression patterns, and markedly downregulate important vaccine candidate genes such as cell surface antigens (Daily *et al.* 2005; Siau *et al.* 2008). The natural host environment, which includes genotypic and immunological cues, may therefore strongly affect the transcriptional responses of malaria parasites. To obtain representative transcriptional information from *Plasmodium* parasites, it is therefore valuable to study natural host systems.

The molecular mechanisms that enable successful invasion and establishment of malaria parasites in hosts other than mice and primates are unfortunately poorly known. Which genes are conserved across *Plasmodium* and how do virulence, immune evasion, and host-specificity vary in species infecting non-mammalian animals? To investigate these questions, it will be necessary to assemble and characterize genome-wide expression information from malaria parasites across their host phylogenetic range. With recent developments of high-throughput sequencing techniques, it has now become possible to generate genomic sequences from non-model parasites (Martinsen & Perkins 2013). Dual RNA-sequencing of hosts and their parasites opens up possibilities of simultaneously studying host-parasite interactions and describing transcriptome expression in both actors. Assembling novel parasite sequences *de novo* is, however, a very difficult task. Without a reference genome, transcripts from the host and/or other sources of contamination may remain after annotation and can influence downstream analyses. Meticulous filtering of assemblies using bioinformatics is therefore crucial to avoid erroneous conclusions (see e.g. Koutsovoulos *et al.* 2016). Nevertheless, successfully constructing parasite transcriptome data of high-quality from different hosts will provide us with valuable insights into the hidden biology of *Plasmodium.*

Malaria parasites that infect birds provide an excellent opportunity for studying transcriptional parasite responses due to their enormous diversity and large variation in host specificity and virulence (Bensch *et al.* 2004; Križanauskienė *et al.* 2006; Lachish *et al.* 2011). They are closely related to mammalian *Plasmodium*, but it was only recently established that rodent and primate malaria parasites are indeed monophyletic in the *Plasmodium* phylogeny (Bensch *et al.* 2016). Some avian *Plasmodium* are extreme host generalists, successfully infecting birds over several orders, while other parasites are host specialists infecting a single species (Pérez-Tris *et al.* 2007; Drovetski *et al.* 2014). The avian malaria system allows for the possibility of capturing wild birds in a natural setting, and evaluating their status of malaria infection. Additionally, recent infection experiments have illustrated the potential to study *Plasmodium* in passerines under controlled conditions in laboratories (Zehtindjiev *etal.* 2008; Palinauskas *et al.* 2008, 2011; Cornet *et al.* 2014; Dimitrov *et al.* 2015; Ellis *et al.* 2015).

In this study, we used a bioinformatic multi-step filtering approach to assemble the blood transcriptome of the avian malaria parasite *Plasmodium ashfordi,* mitochondrial lineage GRW2. This parasite was first associated with various warblers (families Acrocephalidae and Phylloscopidae) (Valkiūnas *et al.* 2007), but has been found since 2016 in 15 different host species of two avian orders (Bensch *et al.* 2009). We used our assembly of *P. ashfordi* to evaluate transcriptome characteristics, genome-wide sequence similarity to other apicomplexans, and searched for genes known to be involved in the *Plasmodium* red blood cell invasion process. We analysed expression levels of parasite genes in three experimentally infected birds during two infection stages, peak and decreasing parasitemia, where the hosts have previously been shown to exhibit different transcriptome responses (Videvall *et al.* 2015). This allowed us for the first time to follow and describe the transcriptome of an avian malaria parasite over time in individual hosts.

## Results

### The *Plasmodium ashfordi* transcriptome assembly

We sequenced blood collected from three experimentally infected Eurasian siskins (*Carduelis spinus*) during peak and decreasing parasitemia (day 21 and 31 postinfection) with Illumina dual RNA-sequencing (see Methods for details). The transcriptome of *P. ashfordi* was assembled into two versions in order to make it transparent and as useful as possible for other researchers. The first assembly version, which we refer to as the annotated assembly, contains the transcripts with annotation information from proteins of Apicomplexa (n = 7 860) (Figure 1; Table S1, Supporting information). The second version that we refer to as the total assembly, also includes the unannotated contigs (n = 4 094) that were strictly filtered to remove contigs from the host (Figure 1), resulting in a total of 11 954 representative transcripts (Table 1). The genomes of *Plasmodium* parasites generally contain around 5-6 000 protein-coding genes (Kersey *et al.* 2016), making it reasonable to assume similar gene numbers for *P. ashfordi.* Building eukaryotic transcriptomes *de novo,* however, naturally yields many more transcripts than there are genes in the genome (Grabherr *et al.* 2011), and so the larger number of transcripts in our assembly is a result of isoform varieties, fragmented contigs, and non-coding RNA.

**Figure 1.**
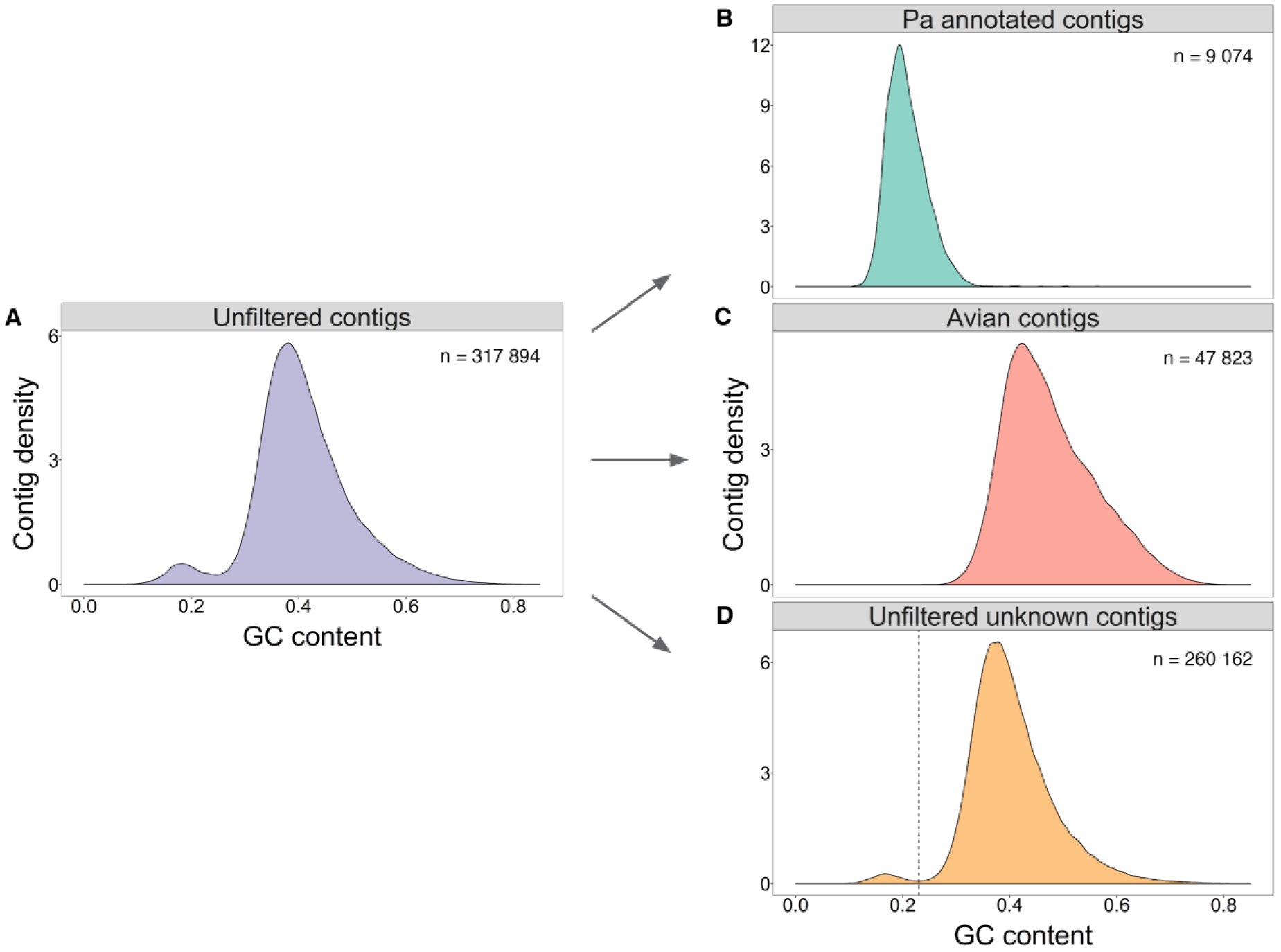
Filtering of host transcripts from the *Plasmodium ashfordi* transcriptome using gene annotation and GC content. Density curves of contig GC content in **(A)** the initial, unfiltered assembly, **(B)** the annotated *P. ashfordi* transcriptome assembly, **(C)** all contigs giving significant blastx matches to birds, and **(D)** all unknown contigs before GC % filtering. The arrows indicate assembly versions before and after initial filtering and cleaning steps. Both the initial, unfiltered assembly and the assembly with unknown, unfiltered contigs display a bimodal distribution, incorporating both avian and malaria parasite transcripts. The dashed straight line in D indicates the 23% GC cut-off where unknown transcripts with lower GC content were extracted, filtered, and later included in the final *P. ashfordi* assembly as unannotated transcripts.

**Table 1.** Assembly statistics of the *Plasmodium ashfordi* transcriptome.

|  | Annotated transcripts | Unannotated transcripts | Total assembly |
| --- | --- | --- | --- |
| Number of contigs | 7 860 | 4 094 | 11 954 |
| Number of bases | 7 316 007 | 1 694 373 | 9 010 380 |
| Contig length min-max (bp) | 200 – 26 773 | 200 – 4 171 | 200 – 26 773 |
| Median contig length (bp) | 681.0 | 321.0 | 498.0 |
| Mean contig length (bp) | 930.8 | 413.9 | 753.8 |
| GC content (%) | 21.22 | 17.26 | 20.48 |

The size of the total transcriptome assembly is 9.0 Mbp and the annotated version of the assembly is 7.3 Mbp (81.20%) (Table 1). We calculated assembly statistics using the transcriptome-specific measurement E90N50, which represents the contig N50 value based on the set of transcripts representing 90% of the expression data, and is preferable over the original N50 when evaluating transcriptome assemblies (Haas 2016). The assembly E90N50 is 1 988 bp and the mean transcript length of the annotated transcripts is 930.8 bp (Table 1; Figure S1, Supporting information). In comparison, the length of coding sequences in the genome of *Haemoproteus tartakovskyi* (bird parasite in a sister genus to *Plasmodium)* is mean = 1 206 and median = 1 809 bp (Bensch *et al.* 2016). The human parasite *Plasmodium falciparum* has a transcriptome with transcripts of median length 1 320 and mean length 2 197 bp (Gardner *et al.* 2002). The longest contig in the *P. ashfordi* assembly, consisting of 26 773 bp, is transcribed from the extremely long ubiquitin transferase gene (AK88_05171), which has a similar transcript length of around 27 400 bp in other *Plasmodium* species. The annotated transcriptome has an exceptionally low mean GC content of 21.22% (Figure 1B), which is even lower than the highly AT-biased transcriptome of *P. falciparum* (23.80%).

### Functional analysis suggests that many host-interaction genes have evolved beyond recognition

To evaluate biological and molecular functions of all annotated transcripts in *P. ashfordi,* we first analysed their associated gene ontology. An analysis of the transcriptome of *P. falciparum* was performed simultaneously to get an appreciation of how the *P. ashfordi* assembly compares functionally to a closely related, well-studied species. Overall, the two transcriptomes displayed highly similar gene ontology patterns (Figures 2A-B; Tables S8-S14, Supporting information). Transcripts primarily belonged to the two major molecular functions: ‘binding’ and ‘catalytic activity’, as well as the biological groups: ‘metabolic process’ and ‘cellular process’. These broad functional classes were also the gene ontology terms where *P. ashfordi* displayed a greater number of transcripts relative *P. falciparum* (Figure 2D). They are likely to contain multiple transcripts per gene due to isoforms, fragmented contigs, or possible gene duplications. The categories ‘receptor activity’, ‘ cell adhesion’, and ‘ multi-organism process’ were in contrast almost exclusively occupied by transcripts from *P. falciparum.* Interestingly, these categories predominately relate to the interaction with host cells and host defences (Figure 2C), which are known to contain genes showing strong evidence for positive selection in *Plasmodium* (Jeffares *et al.* 2007). It is highly likely that many *P. ashfordi* transcripts that belong to these host interaction processes have diverged sufficiently from mammalian parasite genes to escape annotation, but are expressed and present in the unannotated portion of the transcriptome assembly.

**Figure 2.**
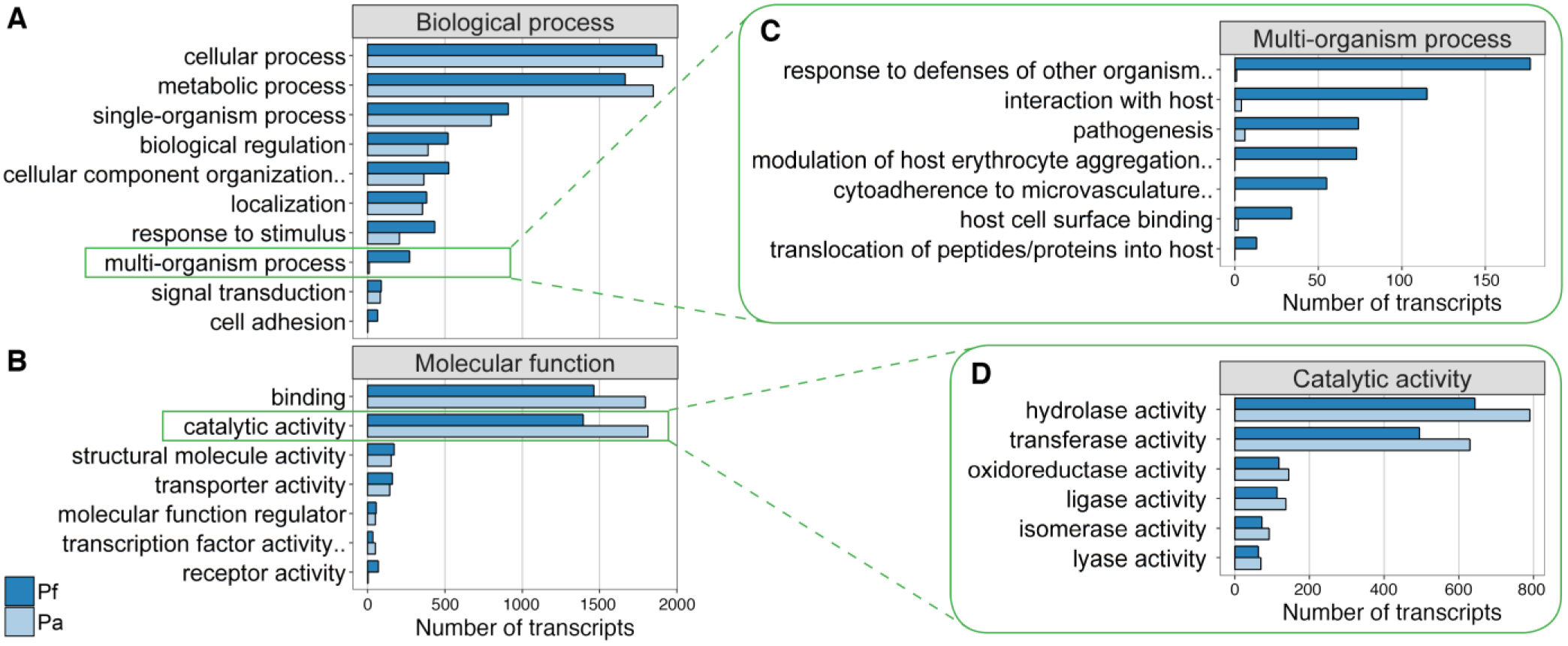
Gene ontology terms of transcripts in the *Plasmodium ashfordi* transcriptome assembly (Pa; light blue) compared to the transcriptome of the human parasite *P. falciparum* (Pf; dark blue). Information of gene ontology was successfully retrieved for 4 346 annotated *P. ashfordi* transcripts and 3 751 *P. falciparum* transcripts. Gene ontology terms containing at least 50 transcripts in one of the species under the two major categories **(A)** ‘biological process’ and **(B)** ‘molecular function’ are shown. Details of underlying child terms are given for two categories: **(C)** ‘multi-organism process’ and **(D)** ‘catalytic activity’, where *P. ashfordi* displays fewer and more transcripts, respectively, relative *P. falciparum.* Gene ontology terms containing at least 10 transcripts in one of the species are shown in C-D. Names of terms ending with dots have been shortened in the figure due to space limitations. For a complete list of terms, see Tables S8-S14 (Supporting information).

We explored the metabolic processes of *P. ashfordi* by looking into the child terms of this gene ontology. All metabolic categories of *P. ashfordi* contained similar or slightly more transcripts compared to *P. falciparum*, except the ‘catabolic process’ where *P. ashfordi* had fewer transcripts (Table S12, Supporting information). An investigation of the broad gene ontology category ‘kinase activity’ resulted in a total of 235 transcripts (Table S1, Supporting information). The kinase multigene family FIKK has drastically expanded in the genomes of *P. falciparum* and *P. reichenowi* (Otto *et al.* 2014b), but only exists as one copy in the rodent malaria parasites. We used the proteins in the FIKK family from the genomes of *P. falciparum* (n = 21) and *P. yoelii* (n = 1) to search for matches in our assembly. All 22 protein sequences produced significant blast matches (e-value < 1e-5) against a single *P. ashfordi* transcript (TR71526|c0_g2_i1), further indicating that the FIKK gene expansion in primate malaria parasites likely happened after the split from both avian and rodent *Plasmodium.* Complete results of all gene ontology analyses of *P. ashfordi* and *P. falciparum* can be found in Tables S8-S14 (Supporting information).

### Gene expression is similar across different stages of infection

Next, we analysed expression levels of the *P. ashfordi* transcripts within individual hosts across the two parasitemia stages. We accounted for differences in parasitemia levels between hosts and time points, and any variation in sequencing depth between samples, by normalizing overall expression values according to the DESeq method (Anders & Huber 2010). We found that the parasites displayed very similar gene expression patterns during peak and decreasing parasitemia stages (Figure 3A-E). No genes were significantly differentially expressed between the time points (q-value > 0.99), and the correlation in gene expression was extremely high (Pearson’s product-moment correlation = 0.9983, t = 1 905.2, df = 11 952, p-value < 2.2e-16) (Figure 3A; Table S2, Supporting information). Annotated transcripts showing the highest expression fold change (non-significantly) between the two parasitemia stages were derived from the following genes (in order of most observed change): rho-GTPase-activating protein 1, 40S ribosomal protein S3a, two uncharacterized proteins, TATA-box-binding protein, heat shock protein 90, and C50 peptidase (Figure 3D; Table S2, Supporting information).

**Figure 3.**
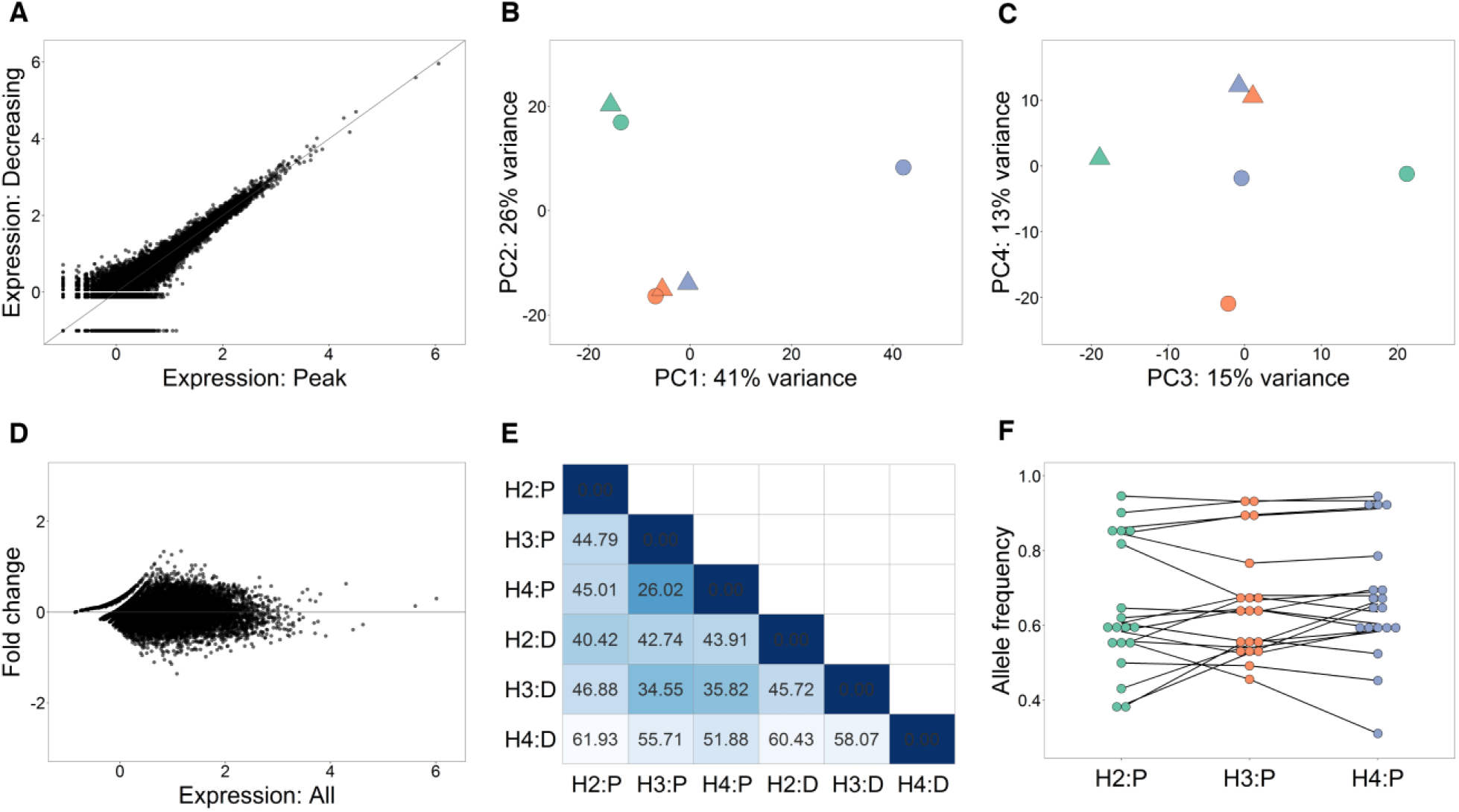
Gene expression patterns of *Plasmodium ashfordi* in individual hosts during two infection stages. **(A)** Scatter plot displaying expression levels of all transcripts in the *P. ashfordi* transcriptome (n = 11 954). The axes show log-transformed normalized mean expression values + 0.1 during peak parasitemia stage (x-axis; n = 3) and decreasing parasitemia stage (y-axis; n = 3). **(B-C)** Principal component analysis (PCA) plots show clustering of samples based on variation in regularized log-transformed normalized gene expression. **(B)** shows principal component 1 and 2, and **(C)** shows principal component 3 and 4. Colours of parasite transcriptomes illustrate which host they are sampled from: host 2 = green, host 3 = orange, host 4 = purple. Triangles and circles, respectively, indicate parasites sampled during peak and decreasing parasitemia stages. **(D)** MA plot showing log-transformed normalized expression values + 0.1 in *P. ashfordi* averaged over all samples (x-axis; n = 6) and shrunken log2 expression fold changes between the parasitemia stages (y-axis). **(E)** Heatmap portraying Euclidian distance measures between parasite expression profiles in different hosts during the two time points. Lighter colour signifies greater distance. H = host, P = peak parasitemia, and D = decreasing parasitemia. The distances between parasite transcriptomes in the decreasing parasitemia stage to any transcriptome sampled during peak parasitemia are shortest within the same host individual. **(F)** Dot plot illustrating allele frequencies of 19 SNPs in the *P. ashfordi* transcriptome from three hosts during peak parasitemia stage. Black lines connect the same SNP for comparisons between hosts. All SNPs contained identical alleles in the parasites from the different hosts and displayed no differences in allele frequency between the host individuals (p-value = 0.432).

### Gene expression is host-specific

In contrast to the similarities in gene expression between parasitemia stages, the parasite transcriptomes showed much larger differences in expression levels between the different host individuals. A principal component analysis of expression variance clustered parasite samples together within their respective hosts, which showed major similarities in expression profiles (Figure 3B). Samples derived from the same host individual did not separate until the third (15% variance) and fourth (13% variance) principal component dimensions (Figure 3C). The parasite transcriptome from host 4 during decreasing parasitemia showed the largest variation in parasite gene expression among all samples, yet it was still most similar to the transcriptome from the same host during peak parasitemia (Figure 3B; Figure 3E). In fact, all parasite transcriptomes during the decreasing parasitemia stage demonstrated closest distance to the transcriptome sample within the same host ten days earlier (Figure 3E). We further evaluated if specific transcripts contributed to the differences in parasite gene expression levels between individual hosts by performing a likelihood ratio test over all host individuals while controlling for parasitemia stage (time point). This resulted in 28 significant *P. ashfordi* transcripts (q-value < 0.1) displaying very high expression variation between hosts (Figure 4; Table S3, Supporting information). The most significant transcripts were derived from the genes cytochrome c oxidase subunit 1, 70 kd heat shock-like protein, M1 family aminopeptidase, and metabolite/drug transporter (Table S3, Supporting information).

**Figure 4.**
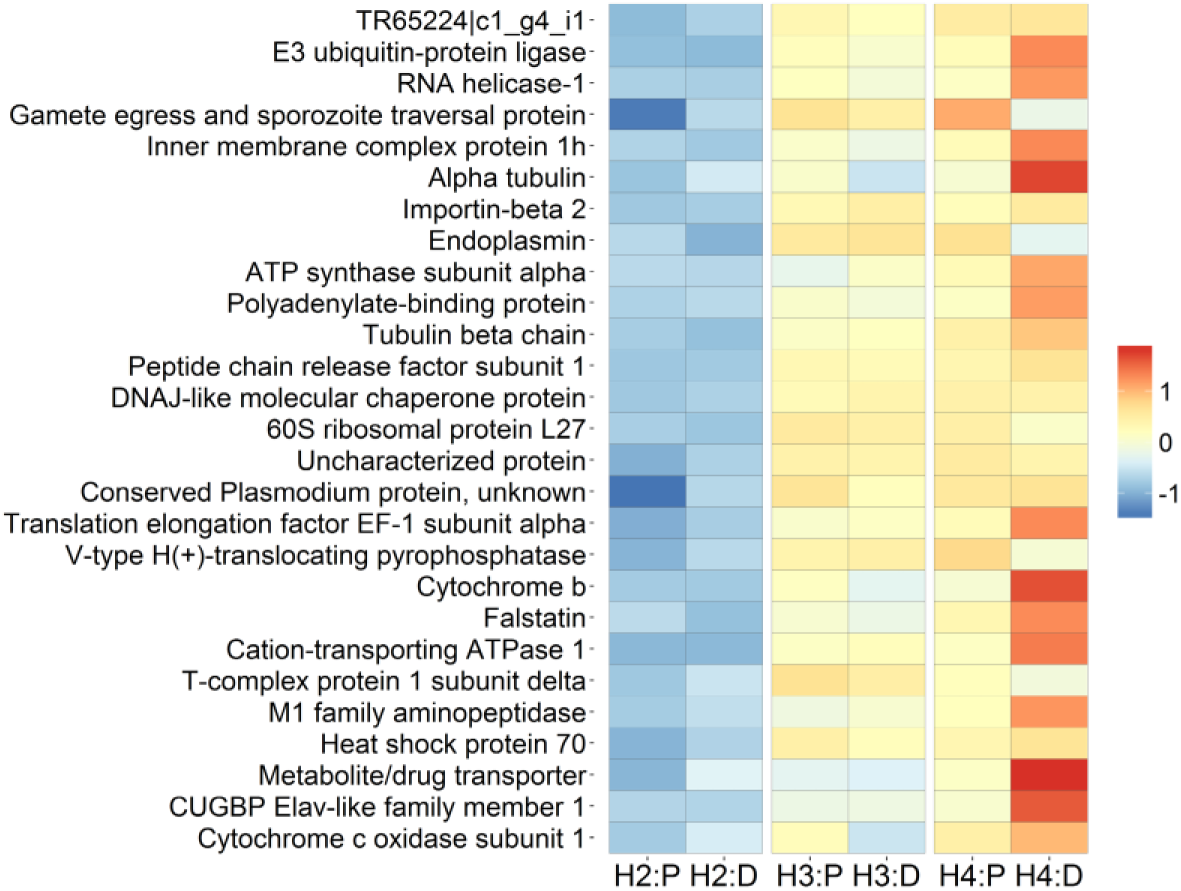
Heatmap of relative expression levels of 28 *Plasmodium ashfordi* transcripts (rows) that were significantly differentially expressed between parasites from different hosts (columns). Warmer colour signifies higher transcript expression, and blue colour indicates lower expression. H = host, P = peak parasitemia, and D = decreasing parasitemia. To compare across genes, expression levels have been normalized with respect to library size, regularized log-transformed, and scaled and centred around zero to give Z-scores.

### *P. ashfordi* is genetically identical in all hosts

In an effort to investigate potential mechanisms behind the host-specific gene expression, we first evaluated if different haplotypes (multiclonality) of the parasite were present and had established differentially in the host individuals. As described in the Methods, all hosts were infected with the same clonal parasite isolate determined by the cytochrome b locus. This was also independently verified in all samples by examining read mapping of mitochondrial transcripts (Figure S5, Supporting information). Because sexual recombination of *Plasmodium* takes place in the mosquito, multiple alleles of nuclear genes could have been present in the parasite strain injected into the birds. A sensitive single nucleotide polymorphism (SNP) analysis over the entire *P. ashfordi* transcriptome found, however, extremely little variation in the parasite. During peak infection, we recovered a total of 10 (unfiltered) SNPs in the 11 954 *P. ashfordi* transcripts from the parasites in host 2, 32 SNPs in host 3, and 46 SNPs in host 4. The variation in number of SNPs called between parasites in different host individuals was due to read coverage, which is directly dependent on parasitemia levels, where host 2 had the lowest, host 3 intermediate, and host 4 highest parasitemia (see Methods). After filtering on read depth, a total of 19 SNPs were identified and used in analyses (Table S4, Supporting information). We discovered that all SNPs (100%) were present in the parasites of all three host individuals, which carried the exact same allele polymorphism (e.g. C/T). To determine if the two alleles were expressed differentially in the hosts, allele frequency was calculated for each SNP in the parasite transcriptome from each host during peak infection. An analysis of the SNPs found no differences in allele frequencies between host individuals (Friedman rank sum test, Friedman chi-squared = 1.68, df = 2, p-value = 0.432), indicating similar allele expression levels in all parasite samples (Figure 3F). The SNPs were also found to be present in the parasites during the decreasing parasitemia stage, but lower coverage during this time point prevents statistical testing.

The 19 SNPs were located inside nine out of 11 954 contigs in the *P. ashfordi* transcriptome, resulting in an allelic occurrence of 0.075%. For a visual example of a contig with three SNPs present, see Figure S6 (Supporting information). Four of the nine transcripts containing SNPs were unannotated, and the others were derived from the genes merozoite surface protein 9 (MSP9), surface protein P113, ubiquitin related protein, multidrug resistance protein, and a conserved *Plasmodium* protein with unknown function (Table S4, Supporting information). We were also able to classify half of the SNPs whether they had any direct effects on the amino acid produced, and found that six SNPs were synonymous and three SNPs were non-synonymous. The three non-synonymous SNPs were located in transcripts from merozoite surface protein 9 (MSP9), ubiquitin related protein, and multidrug resistance protein (Table S4, Supporting information).

The presence of extremely few sequence polymorphisms in the transcriptome of *P. ashfordi* demonstrates that the parasite strain used was not only clonal with respect to the mitochondrial lineage, but homogeneous over the vast majority of the transcriptome. Furthermore, the detection of 19 SNPs that were identical in the parasites from all host individuals verifies that the parasite is genetically identical in all hosts. Intriguingly, it also means that the *P. ashfordi* isolate originally consisted of a minimum of two different haplotypes which differed at nine genes before the experiment, and that this genetic variation managed to survive in the parasite population during the three week long infection, remaining in similar frequency in all hosts.

### Parasite sexual development does not differ between hosts

We further evaluated the possibility that transcriptome host clustering were due to consistent differences relating to the sexual development of *P. ashfordi* during both time points in the host individuals. Lemieux *et al.* (2009) found that differences in *P. falciparum* gene expression between blood samples from children could be partially explained because the parasite expressed different genes during its sexual development (gametocyte) stage. We counted gametocytes in blood slides from the different samples using microscopy (Table S5, Supporting information) and found no differences in gametocyte proportions between host individuals and time points (Pearson’s chi-squared = 1.59, df = 2, p-value = 0.452, mean peak infection = 6.74%, mean decreasing infection = 10.24%). Furthermore, we directly compared the *P. ashfordi* transcripts showing significant expression differences between hosts (n = 28) to a list of sexual development genes exhibiting gametocyte-specific expression patterns in *P. falciparum* (n = 246) (Young *et al.* 2005). Only one of the 28 genes had a match in the sexual development gene set, namely the metabolite/drug transporter MFS1 (Table S3, Supporting information). However, this single match is not greater than expected by chance (hypergeometric test, p-value = 0.372), given probability of a match across all protein-coding genes in the *P. falciparum* genome (n = 5 344). Together, these results show that the host-specific expression pattern in *P. ashfordi* cannot be explained due to differences in sexual development.

### *P. ashfordi* shows sequence similarities to human malaria parasites

Almost all annotated contigs (99.59%; n = 7 828) resulted in a best blast hit against a species within the genus *Plasmodium* (Figure 5A). The remaining contigs had matches against species within the genera of *Eimeria* (n = 12), *Cryptosporidium* (n = 6), *Neospora* (n = 5), *Babesia* (n = 4), *Hammondia* (n = 2), *Ascogregarina* (n = 1), *Theileria* (n = 1), and *Toxoplasma* (n = 1) (Table S6, Supporting information). The great majority (73.59%) of the contig blast matches were proteins originating from primate parasites, while 25.34% matched rodent parasites, and only 0.92% parasites of birds (Figure 5B).

**Figure 5.**
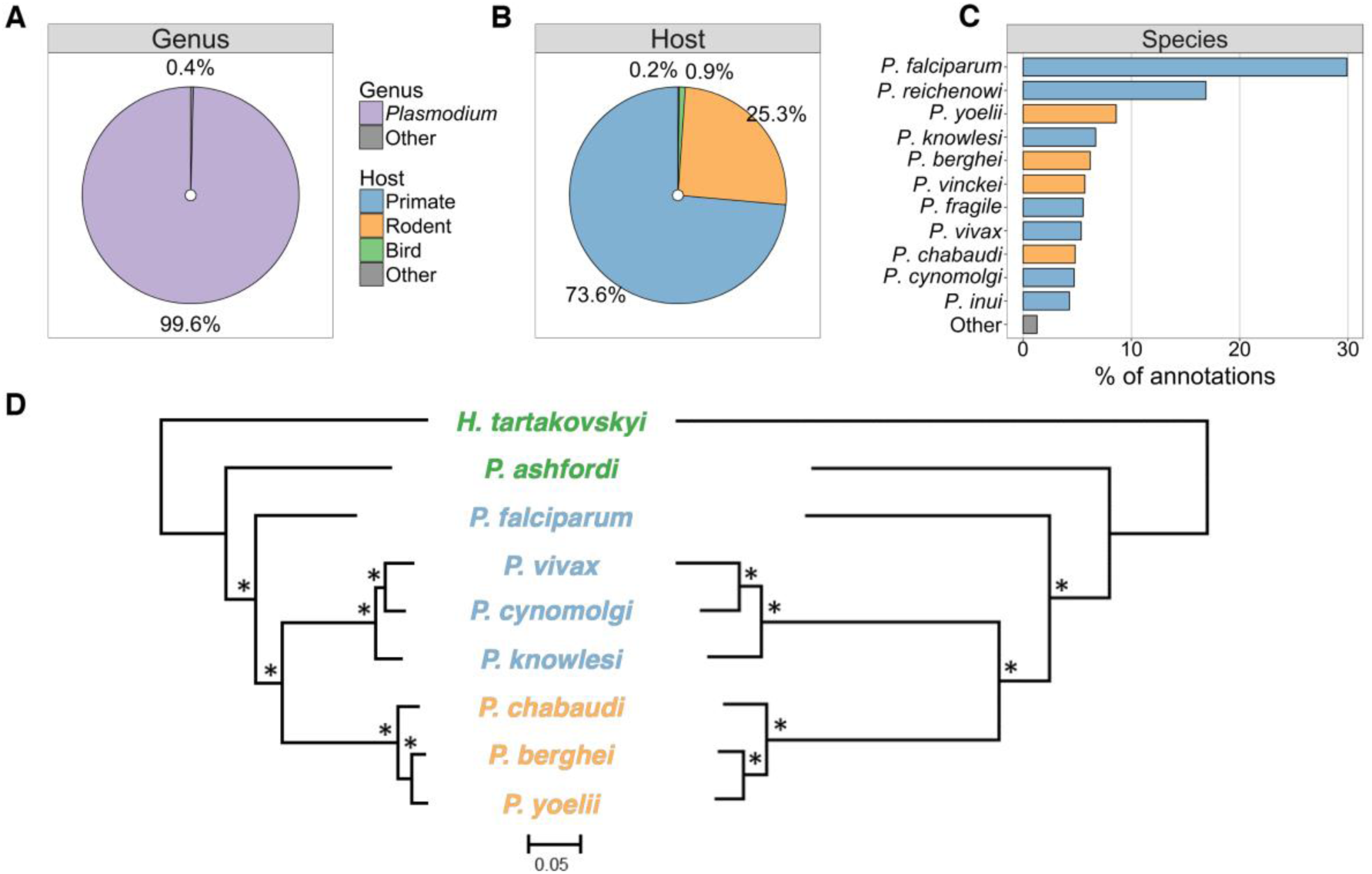
Distribution of apicomplexan parasites presenting best sequence matches to the *Plasmodium ashfordi* transcriptome. **(A)** Pie chart showing the distribution of *P. ashfordi* annotations resulting in significant blast matches to *Plasmodium* species and parasites of other genera. **(B)** Pie chart showing the distribution of annotations resulting in significant matches to parasites infecting primates, rodents, birds, and other hosts. **(C)** Bar plot displaying the proportion of annotated *P. ashfordi* contigs giving significant matches to parasites on a species level. The ‘other’ category contains here all other apicomplexan species, including bird parasites, comprising a total of 20 different species (complete list can be found in Table S6, Supporting information). **(D)** Phylogenetic trees of *Plasmodium* parasites, rooted with the related avian blood parasite *Haemoproteus tartakovskyi.* The left tree is constructed from 599 concatenated genes shared with other species in the Apicomplexa phylum. The right tree is based on 703 concatenated genes that are unique to parasites within these two genera. * = 100% bootstrap support. The phylogenies are derived and adapted with permission from Bensch *et al.* (2016). Same colour scheme as in B applies for C and D, i.e. blue signifies a parasite of primates, orange a parasite of rodents, and green a parasite of birds.

At the species level, most contigs (29.91%) resulted in best blast hit against *P. falciparum,* followed by *P. reichenowi* (16.88%) and *P. yoelii* (8.59%) (Figure 5C). The significant blast matches to bird parasites consisted of the species *Plasmodium gallinaceum* (n = 56), *Eimeria acervulina* (n = 5), *Eimeria tenella* (n = 4), *Eimeria mitis* (n = 3), *Plasmodium relictum* (n = 3), and *Plasmodium lutzi* (n = 1). The contigs giving matches to avian *Plasmodium* were primarily derived from commonly sequenced apicomplexan genes and therefore available in public databases, for example cytochrome c oxidase subunit 1 (COX1; *P. lutzi),* merozoite surface protein 1 (MSP1; *P. relictum),* thrombospondin related anonymous protein (TRAP; *P. relictum),* and cytochrome b (CYTB; *P. gallinaceum)* (Table S1, Supporting information).

The five contigs with highest GC content in the *P. ashfordi* transcriptome (47.7% – 56.4%) all had matches against the avian parasites *Eimeria,* despite them only comprising 0.15% (n = 12) of the total annotation. *Eimeria* species have a very high transcriptome GC content (*E. acervulina*: 55.98%; *E. mitis:* 57.30%), and the *P. ashfordi* transcripts matching this genus consist mostly of ribosomal and transporter genes (Table S1, Supporting information). The *P. ashfordi* contigs with highest expression levels were primarily annotated by uncharacterized protein matches to the rodent parasite *P. yoelii* (Table S7, Supporting information). In fact, the six most highly expressed transcripts that were annotated, all gave significant blast matches to *P. yoelii.* Further investigation revealed that these transcripts are most likely derived from ribosomal RNA.

### Identification of conserved *Plasmodium* invasion genes

Finally, to assess molecularly conserved strategies of *P. ashfordi* compared to mammalian malaria parasites, we searched for annotated genes known to be involved in the red blood cell invasion by *Plasmodium.* (Bozdech *et al.* 2003; Beeson *et al.* 2016). We discovered successfully assembled *P. ashfordi* transcripts from a whole suite of host cell invasion genes (Table 2). This includes for example the genes merozoite surface protein 1 (MSP1), apical membrane antigen 1 (AMA1), merozoite adhesive erythrocytic binding protein (MAEBL), GPI-anchored micronemal antigen (GAMA), and the rhoptry neck binding proteins 2, 4, and 5 (RON2, RON4, and RON5). Interestingly, the *P. ashfordi* RON genes in particular seemed to slightly decrease expression levels in all hosts over the two time points (Figure 6). In general, however, the invasion genes showed a range of expression patterns over time, going in various directions (Figure 6).

**Table 2.** Assembled transcripts of genes involved in *Plasmodium* invasion of red blood cells.

| Gene name | Gene product | Represented by transcripts | Species match (blastx) | Bit score | e-value |
| --- | --- | --- | --- | --- | --- |
| ACT1 | Actin 1 | TR215622\|c0_g1_i1 | *P. vivax* | 758.4 | 5.5E-216 |
| ALP1 | Actin-like protein 1 | TR55613\|c0_g1_i1 | *P. vivax* | 619 | 5.1E-174 |
| AMA-1 | Apical membrane antigen 1 | TR118224\|c0_g1_i1 | *P. gallinaceum* | 624 | 2.1E-175 |
| ARO | Armadillo-domain containing rhoptry protein | TR18124\|c0_g1_i1<br>TR18124\|c0_g2_i1 | *P. knowlesi* | 520<br>501.9 | 1.9E-144<br>6.0E-139 |
| BDP1 | Bromodomain protein 1 | TR66841\|c0_g2_i1<br>TR66841\|c0_g2_i2 | *P. falciparum* | 364.8<br>357.1 | 1.9E-97<br>3.9E-95 |
| FBPA | Fructose-bisphosphate aldolase | TR12911\|c0_g1_i1<br>TR12911\|c0_g2_i1 | *P. cynomolgi*<br>*P. berghei* | 223.8<br>353.2 | 2.5E-55<br>2.7E-94 |
| GAMA | GPI-anchored micronemal antigen | TR16322\|c0_g2_i1 | *P. reichenowi* | 567.4 | 2.5E-158 |
| GAP45 | Glideosome-associated protein 45 | TR34884\|c0_g1_i1 | *P. vivax* | 209.9 | 3.9E-51 |
| GAP50 | Glideosome-associated protein 50 | TR144721\|c0_g1_i1 | *P. knowlesi* | 577.8 | 1.2E-161 |
| MAEBL | Merozoite adhesive erythrocytic binding protein | TR234951\|c0_g1_i1<br>TR99718\|c0_g1_i1<br>TR125315\|c0_g1_i1 | *P. yoelii*<br>*P. gallinaceum*<br>*P. gallinaceum* | 171.8<br>146.4<br>84.3 | 6.3E-40<br>2.4E-32<br>7.2E-14 |
| MCP1 | Merozoite capping protein 1 | TR122100\|c0_g1_i1 | *P. berghei* | 164.5 | 1.7E-37 |
| MSP1 | Merozoite surface protein 1 | TR112241\|c0_g2_i1<br>TR176579\|c0_g1_i1 | *P. relictum* | 368.6<br>134.4 | 4.2E-98<br>1.4E-28 |
| MTIP (MLC1) | Myosin A tail domain interacting protein | TR8937\|c0_g1_i1 | *P. vivax* | 295 | 8.6E-77 |
| MTRAP | Merozoite TRAP-like protein | TR188691\|c0_g2_i1 | *P. vinckei* | 98.6 | 1.3E-17 |
| MYOA | Myosin A | TR198550\|c0_g1_i1 | *Babesia microti* | 199.1 | 1.5E-47 |
| RAMA | Rhoptry-associated membrane antigen | TR144799\|c0_g1_i1 | *P. knowlesi* | 70.1 | 5.0E-09 |
| RALP1 | Rhoptry-associated leucine zipper-like protein 1 | TR7446\|c0_g1_i1 | *P. falciparum* | 102.1 | 1.2E-18 |
| RAP1 | Rhoptry-associated protein 1 | TR115690\|c2_g1_i1 | *P. coatneyi* | 236.1 | 1.3E-58 |
| RAP3 | Rhoptry-associated protein 3 | TR13305\|c0_g1_i1 | *P. falciparum* | 202.6 | 9.9E-49 |
| RBP1 (RH1) | Reticulocyte-binding protein 1 | TR53756\|c1_g1_i1 | *P. falciparum* | 84.7 | 8.0E-13 |
| RBP2 (RH2) | Reticulocyte-binding protein 2 | TR208311\|c0_g1_i1<br>TR66282\|c1_g2_i1 | *P. vivax* | 76.6<br>146.7 | 5.9E-11<br>2.1E-31 |
| RBP2b (RH2b) | Reticulocyte-binding protein 2 homolog B | TR87235\|c1_g1_i1<br>TR35437\|c1_g1_i1 | *P. vivax*<br>*P. falciparum* | 97.8<br>86.7 | 8.7E-18<br>1.9E-14 |
| RHOPH1 | High molecular weight rhoptry protein 1 (CLAG9) | TR7367\|c0_g1_i1<br>TR200056\|c0_g1_i1<br>TR233854\|c0_g1_i1 | *P. reichenowi*<br>*P. reichenowi*<br>*P. falciparum* | 272.3<br>82.8<br>61.6 | 7.9E-70<br>2.3E-13<br>4.8E-07 |
| RHOPH2 | High molecular weight rhoptry protein 2 | TR25858\|c0_g2_i1<br>TR25858\|c0_g3_i1<br>TR175810\|c0_g1_i1 | *P. vivax*<br>*P. fragile*<br>*P. vivax* | 542.3<br>335.9<br>180.6 | 7.8E-151<br>5.7E-89<br>3.4E-42 |
| RHOPH3 | High molecular weight rhoptry protein 3 | TR83052\|c6_g1_i1 | *P. falciparum* | 612.8 | 7.3E-172 |
| RIPR | RH5 interacting protein | TR49727\|c0_g2_i1<br>TR49727\|c0_g2_i2 | *P. reichenowi*<br>*P. fragile* | 502.3<br>299.3 | 6.4E-139<br>1.0E-77 |
| ROM1 | Rhomboid protease 1 | TR178976\|c0_g1_i1 | *P. yoelii* | 102.4 | 2.4E-19 |
| ROM4 | Rhomboid protease 4 | TR92176\|c1_g1_i1 | *P. reichenowi* | 783.5 | 2.1E-223 |
| RON2 | Rhoptry neck protein 2 | TR62454\|c0_g1_i1 | *P. cynomolgi* | 659.8 | 8.5E-186 |
| RON4 | Rhoptry neck protein 4 | TR145809\|c0_g1_i1 | *P. reichenowi* | 349.4 | 8.9E-93 |
| RON5 | Rhoptry neck protein 5 | TR67313\|c1_g1_i1 | *P. reichenowi* | 1105.9 | 0 |
| SUB1 | Subtilisin 1 | TR164664\|c0_g1_i1 | *P. inui* | 636.7 | 3.0E-179 |
| SUB2 | Subtilisin 2 | TR169322\|c0_g1_i1 | *P. inui* | 563.9 | 2.7E-157 |
| TLP | TRAP-like protein | TR153597\|c0_g1_i1 | *P. falciparum* | 440.7 | 3.2E-120 |
| TRAMP | Thrombospondin related apical membrane protein | TR179194\|c0_g1_i1 | *P. falciparum* | 231.1 | 2.1E-57 |
| TRAP | Thrombospondin related anonymous protein | TR16495\|c0_g1_i1 | *P. relictum* | 229.2 | 6.4E-57 |

**Figure 6.**
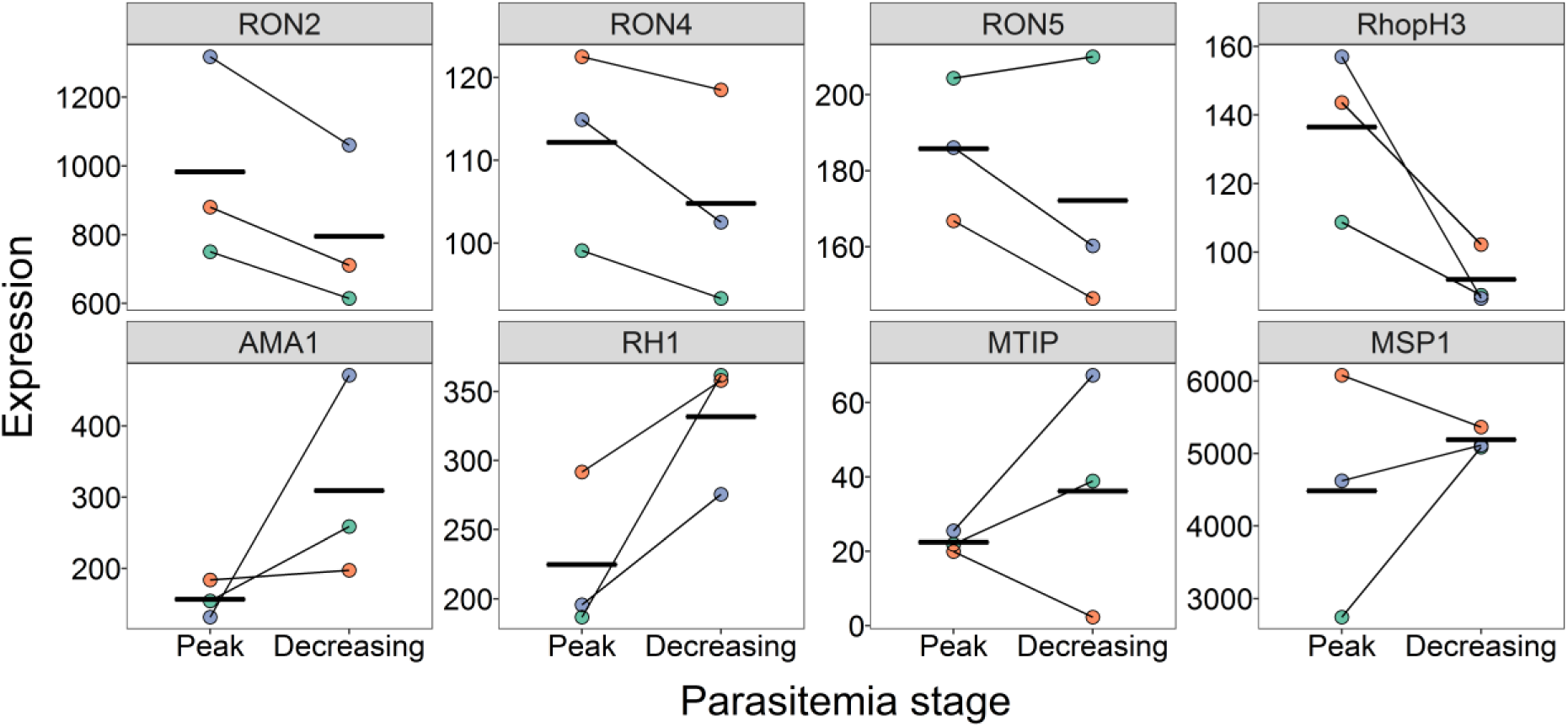
Individual gene expression plots for some of the *Plasmodium ashfordi* transcripts involved in red blood cell invasion. Line plots displaying normalized parasite gene expression in each individual host over the two sampled parasitemia stages. Host 2 is depicted in green, host 3 in orange, and host 4 in purple. Thick horizontal lines indicate mean expression levels in each stage.

All genes known to be involved in the *Plasmodium* motor complex (Opitz & Soldati 2002; Baum *et al.* 2006), driving parasite gliding motion and enabling host cell invasion, were discovered in *P. ashfordi.* These include: actin (ACT1), actin-like protein (ALP1), aldolase (FBPA), myosin A (MyoA), myosin A tail interacting protein (MTIP), glideosome-associated protein 45 and 50 (GAP45 and GAP50), and thrombospondin related anonymous protein (TRAP) (Table 2). We also found the bromodomain protein 1 (BDP1), which has been directly linked to erythrocyte invasion by binding to chromatin at transcriptional start sites of invasion-related genes and controlling their expression (Josling *et al.* 2015).

We found two transcripts matching the low molecular weight rhoptry-associated proteins 1 and 3 (RAP1 and RAP3) that are secreted from the rhoptry organelles during cell invasion. The genomes of human malaria parasites contain a paralog gene called RAP2 as well, whereas rodent malaria parasites contain a single gene copy that is a chimera of RAP2 and RAP3 (RAP2/3) (Counihan *et al.* 2013). The *P. ashfordi* transcript in question (TR13305|c0_g1_i1) matches *P. falciparum* RAP3 better than the rodent parasite version of RAP2/3. The three high molecular weight rhoptry proteins (RhopH1, RhopH2, RhopH3) which bind to the erythrocyte plasma membrane and transfer to the parasitophorous vacuole membrane upon invasion (Vincensini *et al.* 2008; Counihan *et al.* 2013) were all identified in *P. ashfordi.* RhopH1 encompasses the multigene family of cytoadherence linked asexual proteins (CLAGs), present in varying copy number across *Plasmodium.*

Other assembled *P. ashfordi* orthologs of genes involved in host cell invasion were the rhoptry-associated leucine zipper-like protein 1 (RALP1), rhoptry-associated membrane antigen (RAMA), armadillo-domain containing rhoptry protein (ARO), RH5 interacting protein (RIPR), TRAP-like protein (TLP), merozoite TRAP-like protein (MTRAP), thrombospondin related apical membrane protein (TRAMP), subtilisin proteases 1 and 2, (SUB1 and SUB2), and merozoite surface proteins 8 and 9 (MSP8 and MSP9). MTRAP and TRAMP are proteins that belong to the TRAP-family and are released from the microneme organelles during invasion (Green *et al.* 2006; Cowman *et al.* 2012), and the subtilisin proteases SUB1 and SUB2 are heavily involved in the processing and cleavage of immature merozoite antigens, for example MSP1 and AMA1 (Beeson *et al.* 2016). ARO plays a crucial role in positioning the rhoptry organelles within the apical end of the parasite to enable the release of rhoptry-associated molecules (Mueller *et al.* 2013) such as RAMA and RALP1, which then bind to the erythrocyte surface.

We furthermore discovered transcripts of several reticulocyte binding proteins (RBP/RH) thought to be absent in the genomes of avian malaria parasites (Lauron *et al.* 2015). These particular transcripts, together with RAMA, showed much higher e-values than other invasion genes (Table 2), indicating high differentiation between avian and mammalian *Plasmodium* RH genes. Finally, two rhomboid proteases (ROM1 and ROM4) have been linked to host cell invasion in *P. falciparum* via cleavage of transmembrane adhesins (Baker *et al.* 2006; Santos *et al.* 2012). We found both of these genes, together with other rhomboid proteases (ROM2, ROM3, ROM6, ROM8, and ROM10) expressed in *P. ashfordi.* More information about the assembled genes can be found in Table S1 (Supporting information).

## Discussion

In this study, we assembled and characterized a blood-stage transcriptome with quantified gene expression of an avian malaria parasite, *P. ashfordi.* By developing a bioinformatic filtering method capable of dealing with dual RNA-seq data, we effectively removed contigs originating from the host and other sources of contamination in a multistep approach (Figure 1). This resulted in a transcriptome with 7 860 annotated transcripts and an additional 4 094 unannotated transcripts. We discovered that 19 SNPs were not only present, but identical, in the transcriptomes of all six parasite samples from the three hosts, which corroborates that these are indeed true sequence polymorphisms and is an excellent independent verification that all host transcripts have been successfully removed from the assembly (Table S4, Supporting information). The gene expression of *P. ashfordi* displayed strikingly similar patterns during peak and decreasing infection stages and within individual hosts (Figure 3). Furthermore, *P. ashfordi* shows most sequence similarities to the human malaria parasite *P. falciparum* (Figure 5C), but specific genes involved in host interaction and defence seem to be highly differentiated between the two parasites (Figure 2C). Nonetheless, the assembly supports several important erythrocyte invasion genes (Table 2), indicating evolutionary conserved cell invasion strategies across the phylogenetic host range of *Plasmodium* parasites.

### *P. ashfordi* displays host-specific gene expression

Interestingly, and contrary to our expectations, *P. ashfordi* showed highly similar expression profiles inside the same host, despite being sampled ten days apart during two different disease stages. All birds were inoculated with the same malaria strain derived from a single donor bird, and our SNP analysis verified that the genetic composition of *P. ashfordi* is identical in all hosts. The mechanism behind this host-specific expression pattern is unknown, but there is no reason why six independently sampled parasite populations should cluster transcriptionally based on host individual unless expression levels are somehow regulated in response to hosts. The expression pattern can potentially be caused by genotype by genotype interactions between the host and the parasite, modulation of parasite expression by the host, or plasticity of the parasite to different host environments. This result has potentially important implications for our understanding of the evolution of host-parasite interactions, and as a result warrants further research extending the limited sample size to more hosts and more timepoints throughout the infection.

Host genotype by parasite genotype interactions are complicated and not that well documented in malaria parasite systems. Studies with different genotypes of both host and parasite have found effects of host genotype, but not parasite genotype, on factors such as host resistance and parasite virulence (Mackinnon *et al.* 2002; de Roode *et al.* 2004; Grech *et al.* 2006; see also Idaghdour *et al.* 2012). Less is known about the transcriptome responses of malaria parasites to different host individuals. Some studies have found differential gene expression responses of *Plasmodium* to resistant versus susceptible mice strains (see e.g. Lovegrove *et al.* 2006), and Daily *et al.* (2007) discovered host-specific distinct transcriptional states of *P. falciparum* in the blood of Senegalese children. However, a reanalysis of the data by Daily *et al.* (2007) suggested that differences in sexual development of the parasite may have contributed to the transcriptional states (Lemieux *et al.* 2009). With respect to our results, we found no evidence of differences in sexual development of the parasite and our SNP analysis showed that parasite haplotypes did not establish differentially in the host individuals. This suggests that sexual development and haplotype differences had little influence on our results and that the most likely explanation for the host-specific transcriptome patterns is likely to be plasticity in gene expression by *P. ashfordi.*

The expression profiles of *P. ashfordi* did not exhibit any significant differences between peak and decreasing parasitemia stages (day 21 and 31 post-infection). The hosts in our study experienced relatively high parasitemia levels during the decreasing parasitemia stage as well (see Methods), so it is possible that these specific time points do not provide very different environmental host cues to the parasites. However, the concurrent transcriptomes of the avian hosts (analysed in Videvall *et al.* 2015) displayed large differences in gene expression between these two parasitemia stages, notably with a reduced immune response during decreasing parasitemia. This is important because it appears that *P. ashfordi* does not adjust its gene expression in response to the decreasing immune levels of the hosts, but instead conforms to the specific environment of individual hosts.

*P. falciparum* evades the human immune defence via intracellularity, clonal antigenic variation (Guizetti & Scherf 2013), transcriptional antigenic switches (Recker *et al.* 2011), splenic avoidance by erythrocytic adherence to the endothelium (Craig & Scherf 2001) and sequestration in organ microvasculature (Silamut *et al.* 1999), erythrocytic rosetting (Niang *et al.* 2014), host immunosuppression (Hisaeda *et al.* 2004), and manipulation of host gene expression. It is possible that *P. ashfordi* shares several of these evasion strategies with *P. falciparum,* although this remains unknown. One example of immune evasion by manipulation of host gene expression is the parasite gene macrophage migration inhibitory factor homologue (MIF), which contributes to *Plasmodium* parasites’ ability to modulate the host immune response by molecular mimicry (Cordery *et al.* 2007). This gene was discovered transcribed in *P. ashfordi* as well (TR2046|c0_g1_i1), suggesting that a similar immune manipulation strategy is possible (Table S1, Supporting information).

### Similarities to *P. falciparum* and other malaria parasites

The majority of all annotated contigs (73.59%) resulted in a best blast hit against primate parasites (Figure 5B). Curiously, the human malaria parasite *P. falciparum* comprised the majority of all matches with almost a third of the transcripts (29.91%) (Figure 5C). This is likely because *P. falciparum* currently constitutes the organism with most sequence similarities to *P. ashfordi* based on publically available sequences (Bensch *et al.* 2016). The chimpanzee parasite *P. reichenowi* had the second most blast matches to *P. ashfordi* (Figure 5C), and it is the closest living relative to *P. falciparum* based on current genomic data (Otto *et al.* 2014b). Furthermore, both *P. falciparum* and *P. ashfordi* share the genome characteristics of being extremely AT-biased, with *P. ashfordi* reaching a remarkably low transcriptomic GC content of 21.22% (Table 1) compared to the already AT-rich *P. falciparum,* which has a transcriptomic GC content of 23.80%. Lastly, because of its role in human disease, *P. falciparum* is the most sequenced *Plasmodium* species (172 official strains in the NCBI taxonomy database as of May 2016) (Gardner *et al.* 2002), resulting in the greatest opportunity for transcript sequences to find significant blast matches.

Less than one percent of all contigs resulted in a blast hit against avian parasites (0.92%). This is due to the fact that almost no genomic resources are available for avian *Plasmodium.* Despite their enormous diversity, world-wide distribution, and harmful effects on susceptible bird populations, genomic studies of avian malaria parasites have been largely non-existent until now. The genome of *P. gallinaceum,* the malaria parasite of chickens, has been sequenced but not published and we were therefore not able to use it in our analyses. A transcriptome assembly of *P. gallinaceum* is available for download (Lauron *et al.* 2014, 2015), although still contains a large proportion of contigs matching birds, making comparisons with *P. ashfordi* difficult (see Figure S4, Supporting information). Dual RNA-sequencing of a more distantly related apicomplexan parasite, *Leucocytozoon buteonis,* and its buzzard host was recently described (Pauli *et al.* 2015), though no publically available transcriptome exists. Finally, both 454 RNA-sequencing and Illumina genome sequencing of the generalist avian malaria parasite *P. relictum* (lineage SGS1) have been performed (Hellgren *et al.* 2013; Bensch *et al.* 2014; Lutz *et al.* 2016), but the extremely low sequence coverage in both cases does unfortunately not allow for assembly nor any genomic analyses. We hope that future sequencing of these avian parasites will enable genome-wide comparisons.

The lack of genome-wide sequence data from malaria parasites of hosts other than mice and primates means that little is known about which genes across *Plasmodium* are conserved and which that are unique. Our gene ontology results confirm that the transcriptomes of *P. ashfordi* and *P. falciparum* overall are functionally similar (Figure 2A-B), though specific genes involved in host interaction and receptor binding could not be directly located in the *P. ashfordi* assembly (Figure 2C). The transcriptome of *P. falciparum* is certainly more complete because it is based on the genome sequence (Gardner *et al.* 2002) and therefore includes genes expressed from the entire life cycle. As a result, any *P. falciparum* genes not found in *P. ashfordi,* are either specifically transcribed during certain life stages (e.g. in the mosquito), not present in the genome, or too diverged to be detected with sequence similarity searches. Seeing how this particular group of genes involved in host interaction are under strong positive and diversifying selection pressure in *Plasmodium* (Hall *et al.* 2005; Jeffares *et al.* 2007; Otto *et al.* 2014b), it is likely that many are indeed present in the *P. ashfordi* assembly, but unannotated due to evolutionary divergence.

As a step to investigate genes involved in host interaction in more detail, we searched in the *P. ashfordi* assembly for genes specifically known to be involved in the merozoite invasion of host red blood cells. Previously, only a handful of studies have sequenced candidate invasion genes in avian malaria parasites; these include MAEBL (Martinez *et al.* 2013), AMA1 and RON2 (Lauron *et al.* 2014), MSP1 (Hellgren *et al.* 2013, 2015), RIPR (Lauron *et al.* 2015), and TRAP (Templeton & Kaslow 1997; Farias *et al.* 2012). Due to the evolutionary distance between mammals and saurians, and their inherent blood cell differences (birds/reptiles have erythrocytes with a nucleus and mitochondria while mammalian cells are anucleated), we might expect to find few and highly differentiated gene orthologs. Instead, we discovered a large number of red blood cell invasion genes expressed in *P. ashfordi* (Table 2), indicating that most of these specific invasion genes are conserved across both mammalian and avian *Plasmodium.*

The invasion genes that were most differentiated between birds and mammals were the rhoptry-associated membrane antigen (RAMA) and the reticulocyte binding proteins (RBP/RH), which had diverged almost beyond recognition. These RH genes, together with other erythrocyte binding antigens (EBA), have been assumed to be absent in the genomes of avian malaria parasites (Lauron *et al.* 2015). However, our result suggests that several are not only present, but also transcribed, though with high sequence divergence. It is possible that additional erythrocyte binding proteins are present in the *P. ashfordi* assembly, though ortholog searches for these genes will become complicated if they have evolved under especially strong selection pressure in avian *Plasmodium.*

## Conclusion

In this study we have *de novo* assembled, characterized, and evaluated the blood transcriptome of the avian parasite *Plasmodium ashfordi.* By developing a rigorous bioinformatic multistep approach, the assembly was successfully cleaned of host sequences and contains high numbers of important genes in e.g. red blood cell invasion. We have shown that *P. ashfordi* displays similar expression profiles within individual hosts during two different stages of the infection – but different expression patterns between individual hosts – indicating possible host specific parasite gene regulation. The expression information of all transcripts will assist researchers studying genes involved in e.g. immune evasion, host-specificity, and parasite plasticity. In addition, our results show that the isolate of *P. ashfordi* used in this experiment originally contained two alleles at nine loci and has managed to maintain this low-level genomic heterozygosity throughout the infection in all host individuals. The results presented here and the associated assembly will help improve our understanding of host-parasite interactions, evolutionary conserved *Plasmodium* strategies, and the phylogenetic relationships between apicomplexans.

## Methods

### Experimental setup

We used four wild-caught juvenile Eurasian siskins (*Carduelis spinus*) in an infection experiment. The experimental procedure was carried out 2012 at the Biological Station of the Zoological Institute of the Russian Academy of Sciences on the Curonian Spit in the Baltic Sea (55° 05′*N*, 20° 44Έ). All details regarding the setup have been outlined in Videvall *et al.* (2015). Three of the birds were inoculated with a single injection of blood containing the erythrocytic stages of *Plasmodium ashfordi,* subgenus *Novyella,* lineage GRW2. For a description of this parasite, see Valkiūnas *et al.* (2007). A control bird (bird 1) was simultaneously inoculated with uninfected blood for the evaluation of host transcription (see Videvall *et al.* 2015).

The parasite strain was originally collected in 2011 from a single common cuckoo (*Cuculus canorus*) that had acquired the infection naturally. It was thereafter multiplied in common crossbills (*Loxia curvirostra*) in the laboratory and deep frozen in liquid nitrogen for storage. One crossbill was subsequently infected with the parasite, and blood from this single bird was used to infect our experimental birds. Crossbills were used as intermediate hosts because they are both susceptible to this strain and have a large enough body size to provide the amount of blood needed for donations. A subinoculation of a freshly prepared mixture containing infected blood from the donor was made into the pectoral muscle of the recipient birds (details of procedure can be found in Palinauskas *et al.* 2008). By using a single donor, we ensured that the same clonal parasite strain and parasite quantity was injected into recipient birds.

All birds were thoroughly screened with both microscopic (Palinauskas *et al.* 2008) and molecular (Hellgren *et al.* 2004) methods before the experiment to make sure they had no prior haemosporidian infection. Blood samples for RNA-sequencing were taken from birds before infection (day 0), during peak parasitemia (day 21 postinfection) and during the decreasing parasitemia stage (day 31 postinfection). The parasitemia intensity varied substantially in infected birds, with bird 2, bird 3, and bird 4 having 24.0%, 48.0%, and 71.3% of their red blood cells infected during peak parasitemia, and later 8.2%, 21.8%, and 33.3%, respectively, during the decreasing parasitemia stage (Videvall *et al.* 2015). The two parasitemia stages are referred to as different ‘infection stages’ in the hosts, but we want to clarify that there is no evidence present suggesting that the parasites have entered a different stage of their life cycle (e.g. tissue merogony). Experimental procedures were approved by the International Research Co-operation Agreement between the Biological Station Rybachy of the Zoological Institute of the Russian Academy of Sciences and Institute of Ecology of Nature Research Centre (25 May 2010). Substantial efforts were made to minimize handling time and potential suffering of birds.

### RNA extraction and sequencing

From six infected samples (three treatment birds at days 21 and 31) and six uninfected samples (three treatment birds at day 0 and one control bird at days 0, 21, and 31), total RNA was extracted from 20 μl whole blood. Detailed extraction procedures can be found in Videvall *et al.* (2015). Total extracted RNA was sent to Beijing Genomics Institute (BGI), China, for RNA quality control, DNAse treatment, rRNA depletion, and amplification using the SMARTer Ultra Low kit (Clontech Laboratories, Inc.). BGI performed library preparation, cDNA synthesis, and paired-end RNA-sequencing using Illumina HiSeq 2000. The blood samples from bird 3 and bird 4 during peak parasitemia were sequenced by BGI an additional time in order to generate more reads from the parasite in preparation for this transcriptome assembly. These resequenced samples were regarded and handled as technical replicates. We quality-screened all the demultiplexed RNA-seq reads using FastQC (v. 0.10.1) (http://www.bioinformatics.babraham.ac.uk/projects/fastqc/).

### *De novo* transcriptome assembly

Quality-filtered RNA-seq reads from all six infected bird samples together with the two re-sequenced samples were used in a *de novo* assembly. This was performed using the transcriptome assembler Trinity (v. 2.0.6) (Grabherr *et al.* 2011) with 489 million reads. Mapping of reads to available genomes of human malaria parasites were employed simultaneously, but unfortunately these attempts yielded few hits due to the evolutionary distance between avian *Plasmodium* and human *Plasmodium,* so we continued annotating the assembly using blast searches.

The assembled transcripts were blasted against the NCBI non-redundant protein database using the program DIAMOND BLASTX (v. 0.7.9) (Altschul *et al.* 1990; Buchfink *et al.* 2014) with sensitive alignment mode. A total of 47 823 contigs generated significant hits against avian species (Figure 1C). A large number of contigs (n = 260 162) did not produce any significant (e-value < 1e-5) blastx hits (Figure 1D). This is because 1) the host species is a non-model organism without a genome available, leading to a large number of host contigs without blast hits, 2) contigs might not necessarily contain coding sequences, but can be derived from noncoding RNAs, etc. and will therefore not match protein sequences, 3) short, fragmented contigs may not yield sufficient blast hit significance, and 4) an extreme underrepresentation of protein sequences from avian *Plasmodium* species in the NCBI nr database will not result in any significant blast hits to genes unique in avian malaria parasites.

We strictly filtered the initial assembly by only retaining a total of 9 015 transcripts (isoforms) that produced significant blast matches against proteins from species in the Apicomplexa phylum. A previous assembly using an earlier version of Trinity (v. r20140413p1) (Grabherr *et al.* 2011) performed better when it came to assembling the longest contigs (> 6 kbp). Different versions of assembler software may construct de Bruijn graphs somewhat differently, which is why it can be a good idea to make several assemblies and later combine parts of them (Brian Haas, personal communication). The previous assembly had been blasted and screened for Apicomplexa in exactly the same way as described above. In order not to lose these important full-length transcripts, we therefore included the longest contigs from the previous assembly that had 1) not assembled correctly in the current assembly, and 2) significant blastx hits against Apicomplexa (n = 10), resulting in a total of 9 025 transcripts. The fact that these contigs contained similar sequences already present in the assembly was dealt with through downstream clustering of the sequences.

### Transcriptome cleaning and filtering

Some contigs in the annotated assembly contained poly-A tails which complicated downstream analyses and resulted in biased mapping estimates. We therefore removed all poly-A/T tails using PRINSEQ (v. 0.20.4) (Schmieder & Edwards 2011) with a minimum prerequisite of 12 continuous A’s or T’s in the 5’ or 3’ end of the contigs. A total of 106 202 bases (1.18%) was trimmed and the mean transcript length was reduced from 995.28 to 983.52 bases.

The unknown transcripts that failed to produce significant hits to any organism during the blastx run (n = 260 162) were subsequently cleaned using the following procedure. First, we trimmed them for poly-A tails, resulting in a total of 455 331 bases removed, and a slight decrease of the mean length of the unknown sequences from 555.27 nt before trimming to 553.52 nt after trimming. The majority of these unknown transcripts came from host mRNA, but their GC content displayed a clear bimodal distribution (Figure 1D), where the contigs with very low GC were strongly suspected to originate from the parasite. To avoid any host contigs, we strictly filtered the unknown transcripts to only include sequences with a mean GC content lower than 23% (n = 4 624). This threshold was based on the Apicomplexa-matching transcripts (mean GC = 21.22%), the contigs matching birds (class: Aves; n = 47 823; mean GC = 47.65%), and the bird contig with the absolute lowest GC content (GC = 23.48%) (Figure 1C).

### Transcriptome clustering, further filtering, and validation

To reduce redundancy of multiple isoforms and transcripts derived from the same gene, we first merged together the annotated and the unknown transcripts with a GC content < 23% (n = 13 649). We then clustered these sequences together in order to retain most transcripts but group the highly similar ones based on 97% sequence similarity and a k-mer size of 10, using CD-HIT-EST (v. 4.6) (Li & Godzik 2006). The most representative (longest) contig in every cluster was selected to represent those transcripts, resulting in 12 266 contigs/clusters. We further filtered all the short sequences (<200 bases), to obtain a set of 12 182 representative transcripts.

A second blast filtering step with the trimmed representative contigs against the Refseq genomic database was then employed using BLASTN+ (v. 2.2.29) (Altschul *et al.* 1990; Camacho *et al.* 2009) to identify some ambiguous contigs suspected to contain non-coding RNA bird sequences. We removed all contigs that gave significant matches (e-value < 1e-6) against all animals (kingdom: Metazoa), so we could be confident that the assembly only consisted of true parasite transcripts. This last filtering step removed 228 contigs.

The unannotated transcripts (n = 4 094) of the final assembly were further validated to originate from the parasite by using reads from the six uninfected samples of the same hosts sampled before infection and the control bird. A total of 350 318 482 reads (65 bp and 90 bp) from all uninfected samples were mapped to the unannotated transcripts using Bowtie2 (v. 2.2.5) (Langmead & Salzberg 2012), resulting in the alignment of 90 read pairs (0.000051%). This extremely low mapping percentage from the uninfected samples greatly supported our conclusion that these transcripts had indeed been transcribed by *P. ashfordi.* These 4 094 representative transcripts with unknown function are referred to throughout the paper as unannotated transcripts. The resulting final transcriptome assembly consists of 11 954 representative annotated and unannotated transcripts.

### Estimating expression levels

Poly-A tails of 489 million RNA-seq reads from all samples were trimmed as well using PRINSEQ (v. 0.20.4) (Schmieder & Edwards 2011). A minimum prerequisite for trimming were 20 continuous A’s or T’s in the 5’ or 3’ end of each read. Only trimmed reads longer than 40 bp and still in a pair were retained (n = 451 684 626) in order to confidently map high quality reads with good minimum lengths. Bowtie2 (v. 2.2.5) (Langmead & Salzberg 2012) was used to map the trimmed RNA-seq reads of every sample (n = 8) (six biological and two technical replicates) back to the *P. ashfordi* transcriptome consisting of the 11 954 representative sequences. We calculated expression levels using RSEM (v. 1.2.21) (Li & Dewey 2011), which produces expected read counts for every contig.

The counts of the 11 954 transcripts were subsequently analysed inside the R statistical environment (v. 3.2.5) (R Core Team 2015). We tested for expression differences in the malaria parasites between the two time points, and between the hosts, using DESeq2 (v. 1.10.1) (Love *et al.* 2014). The two resequenced samples (technical replicates) of bird 3 and bird 4 during peak parasitemia were handled exactly the same as all other samples, and their respective read count were added to their biological samples, according to the DESeq2 manual. Counts were normalized to account for potential variation in sequencing depth as well as the large differences in number of parasites present in the blood (parasitemia levels). Regularized log transformation of counts was performed in order to represent the data without any prior knowledge of sampling design in the principal component analysis and sample distance calculations. This way of presenting counts without bias is preferred over variance stabilizing of counts when normalization size factors varies greatly between the samples (Love *et al.* 2014), as they naturally do in our data.

### Variant calling and SNP analyses

Sequence variation in the transcriptome assembly was performed according to the Genome Analysis Toolkit (GATK) Best Practices workflow for variant calling in RNA-seq data (v. 2015-12-07). The BAM files for each sample produced by RSEM were sorted, read groups added, and duplicates reads removed using Picard Tools (v. 1.76) (https://broadinstitute.github.io/picard/). With the SplitNCigarReads tool in GATK (v. 3.4-46) (McKenna *et al.* 2010), the mapped reads were further reassigned mapping qualities, cleaned of Ns, and hard-clipped of overhang regions. Variant calling was done in GATK with HaplotypeCaller using ploidy = 1 and optimized parameters recommended for RNA-seq data (see GATK Best Practices). Indels were excluded using the SelectVariants tool, and variants were filtered on quality using the VariantFiltration tool in GATK according to the filter recommendation for RNA-seq data. Next, we used SnpSift in SnpEff (v. 4_3g) (Cingolani *et al.* 2012) to filter variants on depths of a minimum of 20 high-quality, non-duplicated reads. Finally, to calculate allele frequencies of all samples in variant positions called in only some hosts, we ran HaplotypeCaller again in the –ERC BP_RESOLUTION mode and extracted the read depths of the nucleotide positions called previously. Nucleotide positions were filtered if they were positioned in the very end of contigs or had a depth of less than 20 high-quality, non-duplicated reads according to GATK. Allele frequencies were calculated by dividing the number of reads supporting the alternative nucleotide with the total number of reads at each variant site.

### Transcriptome evaluation

Transcriptome statistics such as GC content, contig length, and assembled bases were calculated using Bash scripts and in the R statistical environment (v. 3.2.5) (Pages *et al.* 2015; R Core Team 2015). P-values were corrected for multiple testing with the Benjamini and Hochberg false discovery rate (Benjamini & Hochberg 1995) and corrected values have been labelled as q-values throughout. We calculated the GC content of two *Eimeria* transcriptomes downloaded from ToxoDB (v. 25) (Gajria *et al.* 2007), initially sequenced by Reid *et al.* (2014). Transcriptome E90N50 was calculated using RSEM (v. 1.2.21) and Trinity (v. 2.0.6) (Li & Dewey 2011; Grabherr *et al.* 2011; Haas 2016). Plots were made with the R package ggplot2 (v. 2.2.1) (Wickham 2009). The transcriptome of *Plasmodium falciparum* 3D7 (v. 25) was downloaded from PlasmoDB (Gardner *et al.* 2002; Aurrecoechea *et al.* 2009) and gene ontology information was derived from UniProtKB (Bateman *et al.* 2015). The red blood cell invasion genes were searched for in the transcriptome annotation we produced for *P. ashfordi* (Table S1, Supporting information). Only genes with documented involvement in *Plasmodium* erythrocyte invasion were included in the search.

## Data accessibility

The supplementary tables and figures supporting this article have been uploaded as part of the online supporting information. The sequence reads of both host and parasite have been deposited at the NCBI Sequence Read Archive (SRA) under the accession number PRJNA311546. The assembled *P. ashfordi* transcriptome is available for download as Data S1 and Data S2 (Supporting information), or alternatively at http://mbio-serv2.mbioekol.lu.se/Malavi/Downloads.

## Author contributions

The study design was initially conceived by OH, VP and GV, and further developed together with EV and CKC. The infection experiment was planned by GV and VP, and performed by VP. OH performed the RNA extractions. Assembly and all bioinformatic and statistical analyses were performed by EV. DA advised in the trimming of the contigs and in the mapping of sequence reads. OH, CKC, and EV planned the paper. EV wrote the paper with extensive input from all authors.

## Acknowledgements

We would like to thank Staffan Bensch for stimulating discussions and comments on this paper. We are also grateful to Brian Haas for advice on transcriptome assemblies, and the director of the Biological Station “Rybachy”, Casimir V. Bolshakov, for generously providing facilities for the experimental research. Two anonymous reviewers provided valuable feedback which significantly improved the paper. The assembly and blastn computations were performed on resources provided by SNIC through Uppsala Multidisciplinary Center for Advanced Computational Science (UPPMAX) (Lampa *et al.* 2013) under project b2014134; all other computational analyses were performed by EV on a local machine.

## Supporting Information

**Figure S1.** Length distribution of contigs in the *Plasmodium ashfordi* transcriptome assembly.

**Figure S2.** Multidensity plot showing density of transcripts over log-transformed normalized expression values.

**Figure S3.** Dot plot showing allele frequencies of 19 SNPs in the parasites of three hosts during peak parasitemia.

**Figure S4.** GC content distribution of a transcriptome assembly of the chicken parasite *Plasmodium gallinaceum* (Lauron *et al.* 2015) highlights the difficulties in assembling clean parasite transcriptomes from dual RNA-seq data.

**Figure S5.** Example of transcript sequence homogeneity from the *P. ashfordi* transcript TR213315|c0_g1_i1 derived from the mitochondrial gene cytochrome c oxidase subunit 1 (COX1).

**Figure S6.** Example of transcript sequence heterogeneity showing the presence of three SNPs in the *P. ashfordi* transcript TR73260|c0_g1_i1.

**Table S1**. Information of all annotated *P. ashfordi* transcripts (n = 7 860).

**Table S2**. Normalized expression levels of all *P. ashfordi* transcripts (n = 11 954) in individual hosts during peak and decreasing parasitemia stages.

**Table S3.** *P. ashfordi* transcripts that were significantly differentially expressed between host individuals (n = 28).

**Table S4.** Results of SNPs discovered in the *P. ashfordi* transcriptome (n = 19).

**Table S5.** Gametocyte and meront proportions in the *P. ashfordi* blood smears.

**Table S6.** Species distribution matches from the annotated transcripts of *P. ashfordi.*

**Table S7.** Most highly expressed transcripts in the *P. ashfordi* transcriptome.

**Table S8.** Gene ontology terms with associated numbers of *P. ashfordi* and *P. falciparum* transcripts in the category ‘Biological process’.

**Table S9.** Gene ontology terms with associated numbers of *P. ashfordi* and *P. falciparum* transcripts in the category ‘Molecular function’.

**Table S10.** Gene ontology terms with associated numbers of *P. ashfordi* and *P. falciparum* transcripts in the category ‘ Multi-organism process’.

**Table S11.** Gene ontology terms with associated numbers of *P. ashfordi* and *P. falciparum* transcripts in the category ‘Catalytic activity’.

**Table S12.** Gene ontology terms with associated numbers of *P. ashfordi* and *P. falciparum* transcripts in the category ‘ Metabolic process’.

**Table S13.** Gene ontology terms with associated numbers of *P. ashfordi* and *P. falciparum* transcripts in the category ‘ Cellular process’.

**Table S14.** Gene ontology terms with associated numbers of *P. ashfordi* and *P. falciparum* transcripts in the category ‘Binding’.

**Data S1.** Sequences in the annotated *P. ashfordi* transcriptome assembly (n = 7 860) as a gzipped multi-fasta file.

**Data S2.** Sequences in the total *P. ashfordi* transcriptome assembly (n = 11 954) as a gzipped multi-fasta file.

## References

Altschul SF, Gish W, Miller W, Myers EW, Lipman DJ (1990) Basic local alignment search tool. Journal of Molecular Biology, 215, 403–410.

Anders S, Huber W (2010) Differential expression analysis for sequence count data. Genome Biology, 11, R106.

Aurrecoechea C, Brestelli J, Brunk BP et al. (2009) PlasmoDB: a functional genomic database for malaria parasites. Nucleic Acids Research, 37, D539–D543.

Baker RP, Wijetilaka R, Urban S (2006) Two Plasmodium Rhomboid Proteases Preferentially Cleave Different Adhesins Implicated in All Invasive Stages of Malaria. PLOS Pathogens, 2, e113.

Bateman A, Martin MJ, O’Donovan C et al. (2015) UniProt: a hub for protein information. Nucleic Acids Research, 43, D204–D212.

Baum J, Richard D, Healer J et al. (2006) A conserved molecular motor drives cell invasion and gliding motility across malaria life cycle stages and other apicomplexan parasites. Journal of Biological Chemistry, 281, 5197–5208.

Beeson JG, Drew DR, Boyle MJ et al. (2016) Merozoite surface proteins in red blood cell invasion, immunity and vaccines against malaria. FEMS Microbiology Reviews, 40, 343–372.

Benjamini Y, Hochberg Y (1995) Controlling the False Discovery Rate: a Practical and Powerful Approach to Multiple Testing. Journal of the Royal Statistical Society. Series B, 57, 289–300.

Bensch S, Canbäck B, DeBarry JD et al. (2016) The Genome of *Haemoproteus tartakovskyi* and Its Relationship to Human Malaria Parasites. Genome Biology and Evolution, 8, 1361–1373.

Bensch S, Coltman DW, Davis CS et al. (2014) Genomic Resources Notes accepted 1 June 2013-31 July 2013. Molecular Ecology Resources, 14, 218.

Bensch S, Hellgren O, Pérez-Tris J (2009) MalAvi: a public database of malaria parasites and related haemosporidians in avian hosts based on mitochondrial cytochrome b lineages. Molecular Ecology Resources, 9, 1353–1358.

Bensch S, Pérez-Tris J, Waldenström J, Hellgren O (2004) Linkage between nuclear and mitochondrial DNA sequences in avian malaria parasites: multiple cases of cryptic speciation? Evolution, 58, 1617–1621.

Bozdech Z, Llinás M, Pulliam BL et al. (2003) The Transcriptome of the Intraerythrocytic Developmental Cycle of Plasmodium falciparum. PLOS Biology, 1, e5.

Buchfink B, Xie C, Huson DH (2014) Fast and sensitive protein alignment using DIAMOND. Nature Methods, 12, 59–60.

Camacho C, Coulouris G, Avagyan V et al. (2009) BLAST+: architecture and applications. BMC Bioinformatics, 10, 421.

Cingolani P, Platts A, Wang LL et al. (2012) A program for annotating and predicting the effects of single nucleotide polymorphisms, SnpEff. Fly, 6, 80–92.

Cordery DV, Kishore U, Kyes S et al. (2007) Characterization of a Plasmodium falciparum Macrophage-Migration Inhibitory Factor Homologue. The Journal of Infectious Diseases, 195, 905–912.

Cornet S, Bichet C, Larcombe S, Faivre B, Sorci G (2014) Impact of host nutritional status on infection dynamics and parasite virulence in a bird-malaria system. Journal of Animal Ecology, 83, 256–265.

Counihan NA, Kalanon M, Coppel RL, De Koning-Ward TF (2013) Plasmodium rhoptry proteins: why order is important. Trends in Parasitology, 29, 228–236.

Cowman AF, Berry D, Baum J (2012) The cellular and molecular basis for malaria parasite invasion of the human red blood cell. Journal of Cell Biology, 198, 961–971.

Craig A, Scherf A (2001) Molecules on the surface of the Plasmodium falciparum infected erythrocyte and their role in malaria pathogenesis and immune evasion. Molecular and Biochemical Parasitology, 115, 129–143.

Daily JP, Le Roch KG, Sarr O et al. (2005) In vivo transcriptome of Plasmodium falciparum reveals overexpression of transcripts that encode surface proteins. Journal of Infectious Diseases, 191, 1196–1203.

Daily JP, Scanfeld D, Pochet N et al. (2007) Distinct physiological states of Plasmodium falciparum in malaria-infected patients. Nature, 450, 1091–1095.

Dimitrov D, Palinauskas V, Iezhova TA et al. (2015) Plasmodium spp.: An experimental study on vertebrate host susceptibility to avian malaria. Experimental Parasitology, 148, 1–16.

Drovetski SV, Aghayan SA, Mata VA et al. (2014) Does the niche breadth or trade-off hypothesis explain the abundance-occupancy relationship in avian Haemosporidia? Molecular Ecology, 23, 3322–3329.

Ellis VA, Cornet S, Merrill L et al. (2015) Host immune responses to experimental infection of Plasmodium relictum (lineage SGS1) in domestic canaries (Serinus canaria). Parasitology Research, 114, 3627–3636.

Farias ME, Atkinson CT, LaPointe DA, Jarvi SI (2012) Analysis of the trap gene provides evidence for the role of elevation and vector abundance in the genetic diversity of Plasmodium relictum in Hawaii. Malaria Journal, 11, 305.

Gajria B, Bahl A, Brestelli J et al. (2007) ToxoDB: An integrated toxoplasma gondii database resource. Nucleic Acids Research, 36, D553–D556.

Gardner MJ, Hall N, Fung E et al. (2002) Genome sequence of the human malaria parasite Plasmodium falciparum. Nature, 419, 498–511.

Garnham PCC (1966) Malaria parasites and other Haemosporidia. Blackwell Scientific Publications Ltd., Oxford, UK.

Grabherr MG, Haas BJ, Yassour M et al. (2011) Full-length transcriptome assembly from RNA-Seq data without a reference genome. Nature Biotechnology, 29, 644–652.

Grech K, Watt K, Read AF (2006) Host-parasite interactions for virulence and resistance in a malaria model system. Journal of Evolutionary Biology, 19, 1620–1630.

Green JL, Hinds L, Grainger M, Knuepfer E, Holder AA (2006) Plasmodium thrombospondin related apical merozoite protein (PTRAMP) is shed from the surface of merozoites by PfSUB2 upon invasion of erythrocytes. Molecular and Biochemical Parasitology, 150, 114–117.

Guizetti J, Scherf A (2013) Silence, activate, poise and switch! Mechanisms of antigenic variation in Plasmodium falciparum. Cellular Microbiology, 15, 718–726.

Haas B (2016) Transcriptome Contig Nx and ExN50 stats. https://github.com/trinityrnaseq/trinityrnaseq/wiki/Transcriptome-Contig-Nx-and-ExN50-stats.

Hall N, Karras M, Raine JD et al. (2005) A Comprehensive Survey of the Plasmodium Life Cycle by Genomic, Transcriptomic, and Proteomic Analyses. Science, 307, 82–86.

Hellgren O, Atkinson CT, Bensch S et al. (2015) Global phylogeography of the avian malaria pathogen Plasmodium relictum based on MSP1 allelic diversity. Ecography, 38, 842–850.

Hellgren O, Kutzer M, Bensch S, Valkiūnas G, Palinauskas V (2013) Identification and characterization of the merozoite surface protein 1 (msp1) gene in a host-generalist avian malaria parasite, Plasmodium relictum (lineages SGS1 and GRW4) with the use of blood transcriptome. Malaria Journal, 12, 381.

Hellgren O, Waldenström J, Bensch S (2004) A new PCR assay for simultaneous studies of Leucocytozoon, Plasmodium, and Haemoproteus from avian blood. Journal of Parasitology, 90, 797–802.

Hisaeda H, Maekawa Y, Iwakawa D et al. (2004) Escape of malaria parasites from host immunity requires CD4+CD25+ regulatory T cells. Nature Medicine, 10, 29–30.

Idaghdour Y, Quinlan J, Goulet J-P et al. (2012) Evidence for additive and interaction effects of host genotype and infection in malaria. Proceedings of the National Academy of Sciences of the United States of America, 109, 16786–16793.

Jeffares DC, Pain A, Berry A et al. (2007) Genome variation and evolution of the malaria parasite Plasmodium falciparum. Nature Genetics, 39, 120–125.

Josling GA, Petter M, Oehring SC et al. (2015) A Plasmodium Falciparum Bromodomain Protein Regulates Invasion Gene Expression. Cell Host & Microbe, 17, 741–751.

Kersey PJ, Allen JE, Armean I et al. (2016) Ensembl Genomes 2016: More genomes, more complexity. Nucleic Acids Research, 44, D574–D580.

Koutsovoulos G, Kumar S, Laetsch DR et al. (2016) No evidence for extensive horizontal gene transfer in the genome of the tardigrade Hypsibius dujardini. Proceedings of the National Academy of Sciences of the United States of America, 113, 5053–5058.

Križanauskienė A, Hellgren O, Kosarev V et al. (2006) Variation in host specificity between species of avian hemosporidian parasites: evidence from parasite morphology and cytochrome B gene sequences. Journal of Parasitology, 92, 1319–1324.

Lachish S, Knowles SCL, Alves R, Wood MJ, Sheldon BC (2011) Fitness effects of endemic malaria infections in a wild bird population: The importance of ecological structure. Journal of Animal Ecology, 80, 1196–1206.

Lampa S, Dahlö M, Olason P, Hagberg J, Spjuth O (2013) Lessons learned from implementing a national infrastructure in Sweden for storage and analysis of next-generation sequencing data. GigaScience, 2, 9.

Langmead B, Salzberg SL (2012) Fast gapped-read alignment with Bowtie 2. Nature Methods, 9, 357–359.

Lapp SA, Mok S, Zhu L et al. (2015) Plasmodium knowlesi gene expression differs in ex vivo compared to in vitro blood-stage cultures. Malaria Journal, 14, 110.

Lauron EJ, Aw Yeang HX, Taffner SM, Sehgal RNM (2015) De novo assembly and transcriptome analysis of Plasmodium gallinaceum identifies the Rh5 interacting protein (ripr), and reveals a lack of EBL and RH gene family diversification. Malaria Journal, 14, 296.

Lauron EJ, Oakgrove KS, Tell LA et al. (2014) Transcriptome sequencing and analysis of Plasmodium gallinaceum reveals polymorphisms and selection on the apical membrane antigen 1. Malaria Journal, 13, 382.

Lemieux JE, Gomez-Escobar N, Feller A et al. (2009) Statistical estimation of cell-cycle progression and lineage commitment in Plasmodium falciparum reveals a homogeneous pattern of transcription in ex vivo culture. Proceedings of the National Academy of Sciences of the United States of America, 106, 7559–7564.

LeRoux M, Lakshmanan V, Daily JP (2009) Plasmodium falciparum biology: analysis of in vitro versus in vivo growth conditions. Trends in Parasitology, 25, 474–481.

Li B, Dewey CN (2011) RSEM: accurate transcript quantification from RNA-Seq data with or without a reference genome. BMC Bioinformatics, 12, 323.

Li W, Godzik A (2006) Cd-hit: A fast program for clustering and comparing large sets of protein or nucleotide sequences. Bioinformatics, 22, 1658–1659.

Love MI, Huber W, Anders S (2014) Moderated estimation of fold change and dispersion for RNA-seq data with DESeq2. Genome Biology, 15, 550.

Lovegrove FE, Peña-Castillo L, Mohammad N et al. (2006) Simultaneous host and parasite expression profiling identifies tissue-specific transcriptional programs associated with susceptibility or resistance to experimental cerebral malaria. BMC Genomics, 7, 295.

Lutz HL, Marra NJ, Grewe F et al. (2016) Laser capture microdissection microscopy and genome sequencing of the avian malaria parasite, Plasmodium relictum. Parasitology Research, 115, 4503–4510.

Mackinnon M, Gaffney D, Read A (2002) Virulence in rodent malaria: host genotype by parasite genotype interactions. Infection, Genetics and Evolution, 1, 287–296.

Martinez C, Marzec T, Smith CD, Tell LA, Sehgal RNM (2013) Identification and expression of maebl, an erythrocyte-binding gene, in Plasmodium gallinaceum. Parasitology Research, 112, 945–954.

Martinsen ES, Perkins SL (2013) The diversity of Plasmodium and other haemosporidians: The intersection of taxonomy, phylogenetics and genomics. In: Malaria parasites: comparative genomics, evolution and molecular biology, pp. 1–15. Caister Academic Press, Norfolk.

McKenna A, Hanna M, Banks E et al. (2010) The Genome Analysis Toolkit: A MapReduce framework for analyzing next-generation DNA sequencing data. Genome Research, 20, 1297–1303.

Mueller C, Klages N, Jacot D et al. (2013) The Toxoplasma Protein ARO Mediates the Apical Positioning of Rhoptry Organelles, a Prerequisite for Host Cell Invasion. Cell Host & Microbe, 13, 289–301.

Niang M, Bei AK, Madnani KG et al. (2014) STEVOR Is a Plasmodium falciparum Erythrocyte Binding Protein that Mediates Merozoite Invasion and Rosetting. Cell Host & Microbe, 16, 81–93.

Opitz C, Soldati D (2002) “The glideosome”: a dynamic complex powering gliding motion and host cell invasion by Toxoplasma gondii. Molecular Microbiology, 45, 597–604.

Otto TD, Böhme U, Jackson AP et al. (2014a) A comprehensive evaluation of rodent malaria parasite genomes and gene expression. BMC Biology, 12, 86.

Otto TD, Rayner JC, Böhme U et al. (2014b) Genome sequencing of chimpanzee malaria parasites reveals possible pathways of adaptation to human hosts. Nature Communications, 5, 4754.

Otto TD, Wilinski D, Assefa S et al. (2010) New insights into the blood-stage transcriptome of Plasmodium falciparum using RNA-Seq. Molecular Microbiology, 76, 12–24.

Pages H, Aboyoun P, Gentleman R, DebRoy S (2015) Biostrings: String objects representing biological sequences, and matching algorithms. R package version 2.38.4.

Palinauskas V, Valkiūnas G, Bolshakov CV, Bensch S (2008) Plasmodium relictum (lineage P-SGS1): effects on experimentally infected passerine birds. Experimental Parasitology, 120, 372–380.

Palinauskas V, Valkiūnas G, Bolshakov CV, Bensch S (2011) Plasmodium relictum (lineage SGS1) and Plasmodium ashfordi (lineage GRW2): the effects of the co-infection on experimentally infected passerine birds. Experimental Parasitology, 127, 527–533.

Pauli M, Chakarov N, Rupp O et al. (2015) De novo assembly of the dual transcriptomes of a polymorphic raptor species and its malarial parasite. BMC Genomics, 16, 1038.

Pérez-Tris J, Hellgren O, Križanauskienė A et al. (2007) Within-host speciation of malaria parasites. PLOS ONE, 2, e235.

R Core Team (2015) R: A language and environment for statistical computing. R Foundation for Statistical Computing, Vienna, Austria.

Recker M, Buckee CO, Serazin A et al. (2011) Antigenic Variation in Plasmodium falciparum Malaria Involves a Highly Structured Switching Pattern. PLOS Pathogens, 7, e1001306.

Reid AJ, Blake DP, Ansari HR et al. (2014) Genomic analysis of the causative agents of coccidiosis in domestic chickens. Genome Research, 24, 1676–1685.

de Roode JC, Culleton R, Cheesman SJ, Carter R, Read AF (2004) Host heterogeneity is a determinant of competitive exclusion or coexistence in genetically diverse malaria infections. Proceedings of the Royal Society B: Biological Sciences, 271, 1073–1080.

Santos JM, Graindorge A, Soldati-Favre D (2012) New insights into parasite rhomboid proteases. Molecular and Biochemical Parasitology, 182, 27–36.

Schmieder R, Edwards R (2011) Quality control and preprocessing of metagenomic datasets. Bioinformatics, 27, 863–864.

Siau A, Silvie O, Franetich J-F et al. (2008) Temperature Shift and Host Cell Contact Up-Regulate Sporozoite Expression of Plasmodium falciparum Genes Involved in Hepatocyte Infection. PLOS Pathogens, 4, e1000121.

Siegel T, Hon C-C, Zhang Q et al. (2014) Strand-specific RNA-Seq reveals widespread and developmentally regulated transcription of natural antisense transcripts in Plasmodium falciparum. BMC Genomics, 15, 150.

Silamut K, Phu NH, Whitty C et al. (1999) A Quantitative Analysis of the Microvascular Sequestration of Malaria Parasites in the Human Brain. American Journal of Pathology, 155, 395–410.

Spence PJ, Jarra W, Lévy P et al. (2013) Vector transmission regulates immune control of Plasmodium virulence. Nature, 498, 228–231.

Templeton TJ, Kaslow DC (1997) Cloning and cross-species comparison of the thrombospondin-related anonymous protein (TRAP) gene from Plasmodium knowlesi, Plasmodium vivax and Plasmodium gallinaceum. Molecular and Biochemical Parasitology, 84, 13–24.

Valkiūnas G, Zehtindjiev P, Hellgren O et al. (2007) Linkage between mitochondrial cytochrome b lineages and morphospecies of two avian malaria parasites, with a description of Plasmodium (Novyella) ashfordi sp. nov. Parasitology Research, 100, 1311–1322.

Videvall E, Cornwallis CK, Palinauskas V, Valkiūnas G, Hellgren O (2015) The Avian Transcriptome Response to Malaria Infection. Molecular Biology and Evolution, 32, 1255–1267.

Vincensini L, Fall G, Berry L, Blisnick T, Braun Breton C (2008) The RhopH complex is transferred to the host cell cytoplasm following red blood cell invasion by Plasmodium falciparum. Molecular and Biochemical Parasitology, 160, 81–89.

Wickham H (2009) ggplot2: elegant graphics for data analysis. New York: Springer.

Young JA, Fivelman QL, Blair PL et al. (2005) The Plasmodium falciparum sexual development transcriptome: A microarray analysis using ontology-based pattern identification. Molecular and Biochemical Parasitology, 143, 67–79.

Zehtindjiev P, Ilieva M, Westerdahl H et al. (2008) Dynamics of parasitemia of malaria parasites in a naturally and experimentally infected migratory songbird, the great reed warbler Acrocephalus arundinaceus. Experimental Parasitology, 119, 99–110.

